# Loss of Stathmin-2, a hallmark of TDP-43-associated ALS, causes motor neuropathy

**DOI:** 10.1101/2022.03.13.484188

**Authors:** Kelsey L. Krus, Amy Strickland, Yurie Yamada, Laura Devault, Robert E. Schmidt, A. Joseph Bloom, Jeffrey Milbrandt, Aaron DiAntonio

## Abstract

TDP-43 mediates proper Stathmin-2 (STMN2) mRNA splicing, and STMN2 protein is reduced in the spinal cord of most ALS patients. To test the hypothesis that STMN2 loss contributes to ALS pathogenesis, we generated constitutive and conditional STMN2 knockout mice. Constitutive STMN2 loss results in early-onset sensory and motor neuropathy featuring impaired motor behavior and dramatic distal neuromuscular junction (NMJ) denervation of fast-fatigable motor units, which are selectively vulnerable in ALS, without axon or motoneuron degeneration. Selective excision of STMN2 in motoneurons leads to similar NMJ pathology. STMN2 KO heterozygous mice, which better model the partial loss of STMN2 protein found in ALS patients, display a slowly progressive, motor-selective neuropathy with functional deficits and NMJ denervation. Thus, our findings strongly support the hypothesis that STMN2 reduction due to TDP-43 pathology contributes to ALS pathogenesis.

## Introduction

Loss of *STMN2* expression is hypothesized to contribute to the pathogenesis of amyotrophic lateral sclerosis (ALS). ALS is a neurodegenerative disease characterized by progressive loss of motor function, respiratory failure, and death 3-4 years after symptom onset (van Es *et al*., 2017). Almost all ALS cases, both familial and sporadic, have nuclear loss and/or cytoplasmic inclusions of the RNA-binding protein TDP-43 (Transactive Response DNA-binding protein) in motor neurons and other cells (Ling *et al*., 2013). Two recent landmark studies identified *STMN2* mRNA as the most downregulated transcript in human motor neurons upon TDP-43 knockdown or dysfunction and demonstrated that STMN2 protein levels are reduced in human ALS-affected spinal cords (Klim *et al*., 2019; Melamed *et al*., 2019). These findings imply that most ALS patients will have decreased levels of STMN2 protein and raise the question as to whether or not this loss of STMN2 contributes to ALS pathogenesis. While *STMN2* transcript is a major target of TDP-43, TDP-43 also regulates many other transcripts, some of which likely contribute to ALS pathogenesis (Ma *et al*., 2022; Brown *et al*., 2022). Individually, each key target could play a role in creating the overall phenotype that arises from TDP-43 loss of function. Here we generate constitutive and conditional *Stmn2* mouse mutants to test the hypothesis that a reduction in the levels of STMN2 contributes to ALS pathology.

In human neurons, TDP-43 binds to *STMN2* pre-mRNAs and represses the inclusion of a cryptic exon that results in a prematurely polyadenylated *STMN2* transcript. With the loss of TDP-43, truncated transcript levels rise with concomitant loss of the normal transcript, thereby reducing the amount of functional STMN2 protein in the cell (Klim *et al*., 2019; Melamed *et al*., 2019). Importantly, TDP-43 dependent regulation of STMN2 is not observed in mice because cryptic exons are not conserved between mice and humans (Ling *et al*., 2015). Thus, mouse models with TDP-43 dysfunction do not feature a concurrent loss of STMN2 and only partially mimic an ALS phenotype. Moreover, *Stmn2* mutant mice have not yet been described, so the *in vivo* consequences of STMN2 loss are unknown.

While *in vivo* analysis of STMN2 function is lacking, the molecular and cellular functions of STMN2 are well-studied. STMN2 is a nervous-system specific stathmin family member (Anderson &Axel, 1985; Ozon *et al*., 1997) that binds tubulin dimers to regulate microtubule stability and influences growth cone dynamics to regulate axon outgrowth (Riederer *et al*., 1997; Di Paolo *et al*., 1997). In injured neurons, STMN2 is upregulated and localizes in regenerating growth cones (Shin *et al*., 2014). In addition, STMN2 is an axonal maintenance factor—loss of STMN2 promotes degeneration of injured sensory axons, while overexpression of stabilized STMN2 delays axon degeneration after injury (Shin *et al*., 2012). Interestingly, knockdown of the only stathmin ortholog in *Drosophila* motor neurons leads to neuromuscular junction (NMJ) degeneration and motor axon retraction (Graf *et al*., 2011), while knockout of the stathmin ortholog STMN1 in mice leads to a slowly progressive peripheral neuropathy (Liedtke *et al*., 2002), a disease of axon loss. Hence, the function of stathmins as axonal/synaptic maintenance factors is evolutionarily conserved. In addition to its postulated role in ALS, the truncated *STMN2* transcript is increased in frontotemporal dementia (Prudencio *et al*., 2020), and *STMN2* transcript levels are reduced in Alzheimer’s Disease and Parkinson’s Disease (Mathys *et al*., 2019; Wang Q *et al*., 2019). Indeed, viral-mediated knockdown of STMN2 in the mouse substantia nigra results in dopaminergic neuron death and motor deficits (Wang Q *et al*., 2019). Thus, the role of stathmin family members in axon and NMJ maintenance are consistent with the model that *STMN2* is a functionally relevant target of TDP-43 whose loss contributes to ALS pathology and likely other neurodegenerative disorders.

Here we test the hypothesis that *in vivo* STMN2 depletion results in axon and/or NMJ instability and contributes to ALS pathology. We generated *Stmn2* constitutive and conditional knockout mice and find that STMN2 is required for motor and sensory system function. *Stmn2* deletion results in a severe motor and sensory neuropathy with behavioral defects, reduced compound muscle action potentials, NMJ denervation, and intraepidermal nerve fiber loss. The motor neuropathy in the *Stmn2* KO mouse is most prominent in more distal NMJs, and predominantly affects the fast-fatigable motor units as is seen in ALS (Dengler *et al*., 1990). Conditional deletion of *Stmn2* in choline acetyltransferase (ChAT) expressing neurons results in motor defects and NMJ degeneration, indicating a cell-autonomous role for STMN2 in motor neurons. The cryptic splicing observed in ALS patients with TDP-43 pathology results in a partial loss of STMN2, with postmortem ALS patient spinal cords showing low, but variable, levels of STMN2 (Klim *et al*., 2019). To more closely model the partial loss of STMN2 protein observed in ALS patients, we examined *Stmn2*^*+/-*^ heterozygous mice. These mice exhibit a progressive, distal motor neuropathy with no observed sensory defect. Moreover, loss of this single copy of *Stmn2* results in distal NMJ denervation, an early feature of ALS pathology. These findings demonstrate that STMN2 loss is sufficient to replicate some aspects of ALS disease progression and pathology *in vivo*, and strongly support the hypothesis that the reduction of STMN2 protein levels due to TDP-43 dysfunction contributes to ALS pathogenesis.

## Results

### STMN2 promotes microtubule polymerization and axon outgrowth

Previous genetic analyses of STMN2 used overexpression or knockdown models, not bona fide genetic deletion. To investigate the effects of STMN2 loss *in vivo*, we engineered a constitutive knockout (KO) mouse using CRISPR/Cas9-mediated genome editing. The *Stmn2* gene contains 5 exons spanning ∼55 kb. Just over 1.2 kb were deleted in our model including exon 3, which encodes a large part of the tubulin-binding stathmin-like domain of STMN2, and parts of the surrounding introns (Fig. 1A). This deletion was identified by PCR and subsequently confirmed by DNA sequencing. Heterozygous *Stmn2* mutant mice were then bred to produce *Stmn2* knockout (KO) animals. Loss of STMN2 protein was confirmed by Western blot (Fig. 1B).

**Figure 1:**
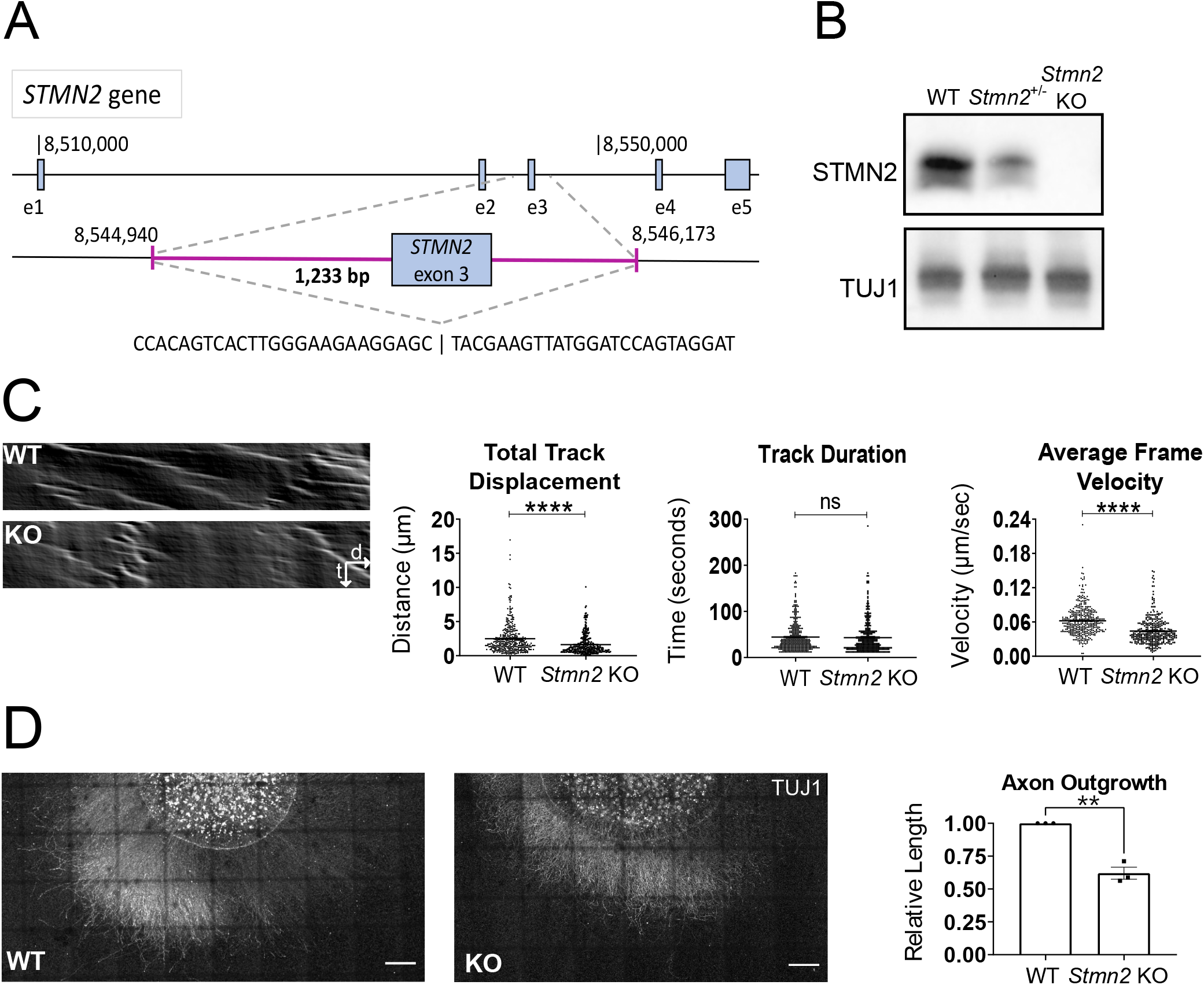
Constitutive *Stmn2* KO results in delayed microtubule polymerization and axon outgrowth. **A)** Schematic of *Stmn2* gene deletion region. Magenta area represents deleted region. **B)** STMN2 protein in brain lysates from 3-month-old mice heterozygous and homozygous for *Stmn2* deletion allele. **C)** Representative kymographs of EB3-mNeonGreen movement from wildtype (WT) and *Stmn2* KO (KO) embryonic DRG neurons. Quantified to the right is total track displacement (µm), track duration (seconds), and average frame velocity (µm/second). **D)** Representative images of spot culture axon length on DIV3. Relative axon length quantified to the right. Scale bar = 500 µm. All data are presented as mean +/-SEM. Statistical significance was determined by Student’s unpaired t-test. (ns: not significant, **p<0.01, ****p<0.0001)

To examine STMN2 loss in individual neurons, we first performed *in vitro* studies. In biochemical assays, STMN2 binds tubulin heterodimers to promote microtubule catastrophe (Riederer *et al*., 1997). In neuronal cultures, it regulates microtubule morphology in growth cones (Morii *et al*., 2006). To assess the effects of STMN2 loss on tubulin polymerization dynamics in a cellular context, we used lentivirus to express a mNeonGreen-tagged EB3 (Chertkova *et al*., 2017), a microtubule plus-end binding protein, in wildtype (WT) and *Stmn2* KO dorsal root ganglion (DRG) neurons. Compared to microtubules in WT neurons, microtubules in *Stmn2* KO neurons show a similar track length but a shorter track displacement, indicating that the rate of tubulin polymerization is slower in the absence of STMN2 (Fig. 1C). This demonstrated defect in tubulin polymerization, along with prior *STMN2* knockdown studies showing decreased axon outgrowth (Morii *et al*., 2006), led us to examine axon outgrowth in our *Stmn2* KO neurons. We cultured embryonic DRG neurons for three days and then measured the length of the axon halo surrounding the cell bodies. We found that axon outgrowth from *Stmn2* KO neurons was on average 35% shorter than the outgrowth from WT neurons (Fig. 1D). Hence, STMN2 is required for normal microtubule polymerization and axon outgrowth but is not essential for either process. Having identified molecular and cellular defects in *Stmn2* KO neurons, we next explored the *in vivo* requirement for STMN2.

### Stmn2 Knockout Mice Display Reduced Perinatal Viability

The *Stmn2* KO mice were first assessed by examining whether STMN2 was required for development. To explore viability, we determined the genotypes of progeny from *Stmn2*^*+/-*^ heterozygous animals one day prior to birth (E18.5) and at three weeks of age. At E18.5, *Stmn2* KO embryos comprised ∼25% of progeny, however by three weeks of age there was a deviation from Mendelian ratios as only 8% were homozygous for the S*tmn2* KO allele (Fig. 2A). While perinatal death is increased, we observed very little premature death after weaning. Indeed, the surviving homozygous *Stmn2* KO mice appear grossly normal, and do not exhibit any obvious gait changes or paralysis up to 1 year of age. Hence, *Stmn2* KO embryos are viable *in utero* but have reduced perinatal fitness.

**Figure 2:**
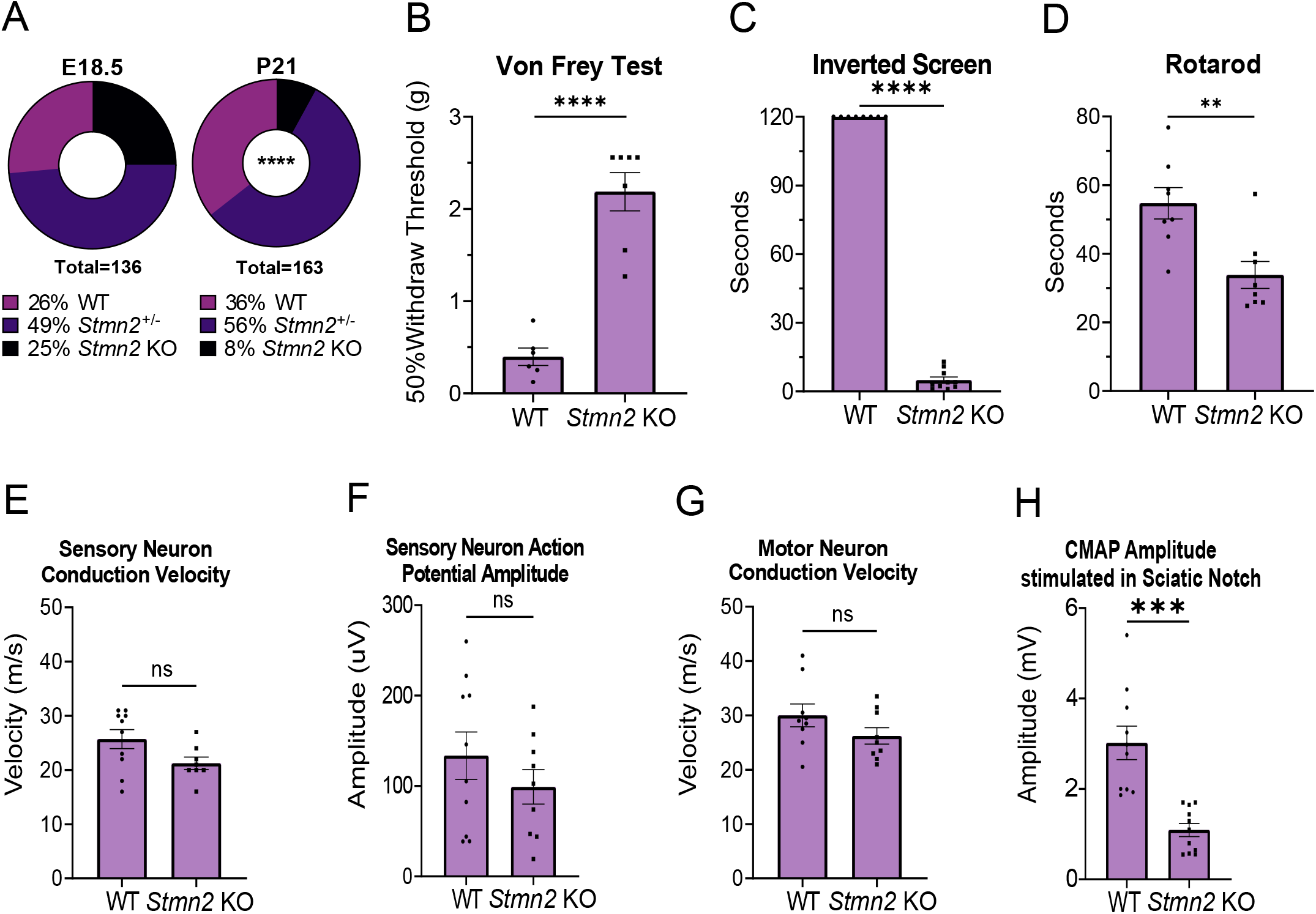
Total STMN2 loss causes perinatal lethality, sensory and motor deficits. **A)** Genotype distribution of E18.5 embryos and P21 mice. Statistical significance was determined by Chi-squared test for goodness of fit. **B)** Average 50% hind paw withdrawal threshold when force (grams) applied, **C)** Latency time (seconds) to fall from an inverted screen (max. 120 sec), and **D)** Length of time (seconds) on an accelerating rotarod before falling, for 3-month-old WT and *Stmn2* KO mice. **E)** Average sensory nerve conduction velocity and **F)** Action potential amplitude for 3-month-old WT and *Stmn2* KO mice. **G)** Average motor neuron conduction velocity and **H)** Compound muscle action potential (CMAP) amplitude stimulated in the sciatic notch for 3-month-old WT and *Stmn2* KO mice. Unless otherwise stated, statistical significance was determined by Student’s unpaired t-test. (ns: not significant, **p<0.01, ***p<0.001, ****p<0.0001).

The defects observed in axon outgrowth *in vitro* led us to hypothesize that *Stmn2* KO embryos might show defects in peripheral nerves. We focused on the phrenic nerve, as loss of diaphragm innervation would be incompatible with life. Analysis of whole-mount preparations stained for the axonal protein neurofilament revealed that innervation of the diaphragm is grossly normal in the *Stmn2* KO embryos. To assess neuromuscular junction (NMJ) development, we imaged postsynaptic acetylcholine receptors (AChR), axons, and synaptic vesicles. We found diaphragm innervation to be very similar in wild-type and *Stmn2* KO embryos, with presynaptic terminals well apposed to postsynaptic AChR clusters, with no significant differences in the numbers of primary and secondary neurites, length of primary branches, or number and size of AChR clusters. *Stmn2* KO animals do have a mild decrease in secondary neurite length compared to WT animals (Supplementary Fig. 1A-B). While other developmental defects may occur in *Stmn2* KO animals that die soon after birth, we conclude that peripheral nerve outgrowth and innervation of the diaphragm are grossly normal and so are not the cause of perinatal lethality.

STMN2 has three close protein orthologs, STMN1, STMN3, and STMN4. We investigated whether any of these proteins were upregulated in surviving three-month old *Stmn2* KO mice to explain why some mice survive to adulthood. We performed Western blot analysis of brain tissue from wild type, *Stmn2* KO, and *Stmn2*^*+/-*^ mice but did not find significant differences in these orthologs in *Stmn2* mutants (Supplementary Fig. 2). Hence, we have no evidence that ortholog compensation explains the viability of the surviving *Stmn2* KO animals.

### Stmn2 Knockout Mice Exhibit Distal Motor and Sensory Neuropathy

If loss of STMN2 contributes to ALS pathology, then we would predict motor defects in the adult *Stmn2* KO animals. To explore the requirement for STMN2 in the adult nervous system, we assayed sensory and motor behavior in three-month-old *Stmn2* KO mice. Although the mice appear healthy with grossly normal behavior, we found they have profoundly impaired mechanosensory perception (assessed by the Von Frey test) and grip strength (assessed by the hanging wire test) (Fig. 2B,C). The mutant mice also display reduced motor coordination on the rotarod, consistent with prior findings using viral-mediated *STMN2* knockdown in the mouse midbrain (Wang *et al*., 2019) (Fig. 2D). We next performed electrophysiological assays of motor and sensory fibers which revealed normal conduction velocities in both sensory and motor nerves in *Stmn2* KO mice, suggesting normal axon myelination (Fig. 2E,G). Similarly, the amplitude of the sensory nerve action potential measured in the tail is normal in *Stmn2* KO mice, indicating that axon loss is minimal in sensory nerves. In contrast, there is a large reduction in the compound motor action potential (CMAP) of *Stmn2* KO mice compared to control animals following stimulation at the sciatic notch and recording at the foot (Fig. 2F,H). Defects in CMAP likely reflect either axon loss or NMJ denervation, consistent with the motor deficits in these mice.

To determine if peripheral axons are lost, we counted myelinated axons in nerve sections. The sciatic, femoral, and sural nerves from *Stmn2* KO mice all have a normal axon density at three and twelve months of age (Fig. 3A, Supplementary Fig. 3A). These represent mixed (sciatic), primarily motor (femoral), and primarily sensory (sural) nerves, indicating that constitutive STMN2 deletion does not cause axon degeneration within the nerve of either fiber type. Consistent with the preservation of motor axons, we observed no decrease in motor neuron number in the lumbar spinal cord of *Stmn2* KO animals, indicating that STMN2 loss does not cause motor neuron death (Supplementary Fig. 3B).

**Figure 3:**
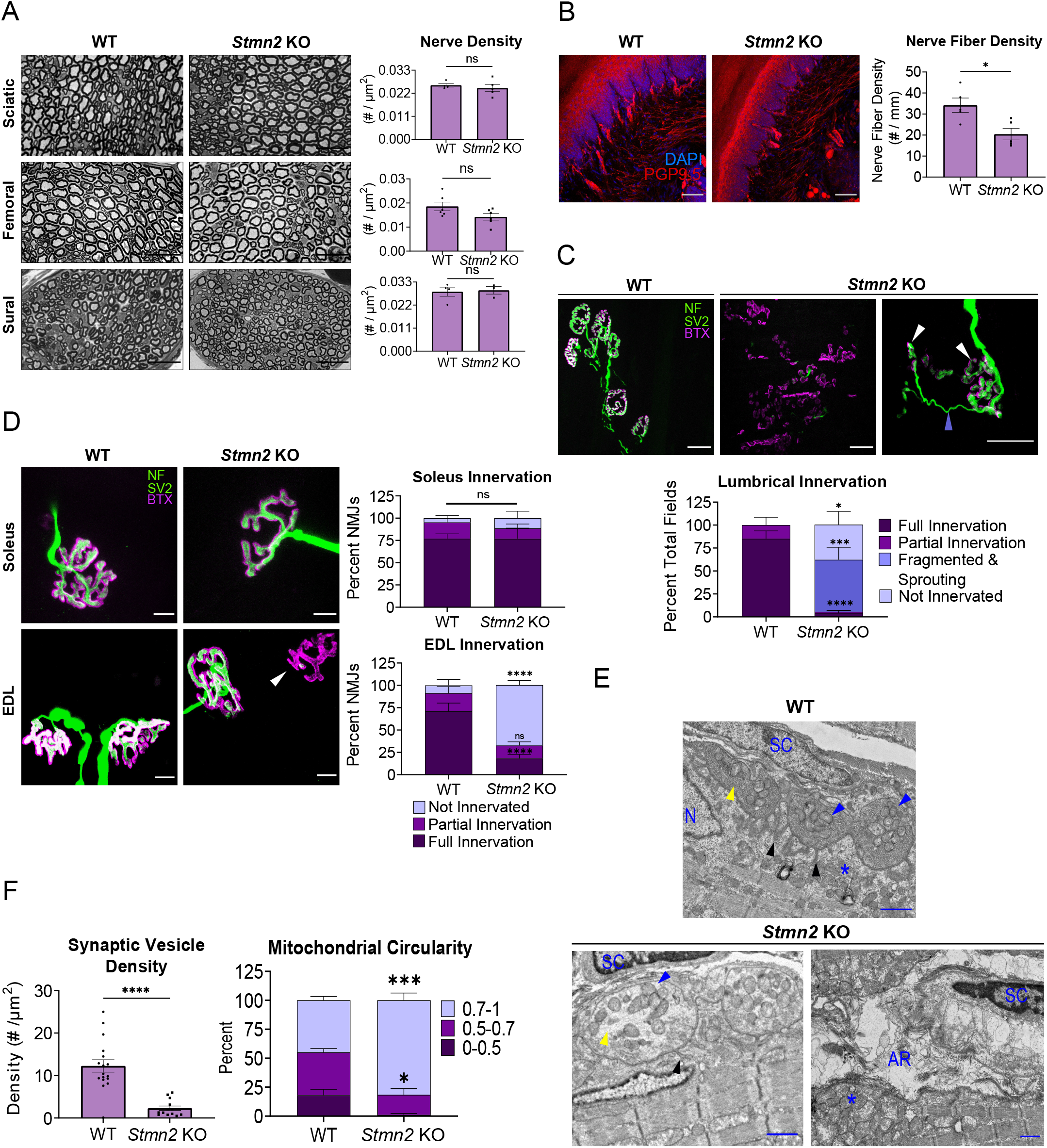
STMN2 deletion results in intraepidermal nerve fiber loss and distal NMJ denervation. **A)** Representative images of sciatic, femoral, and sural nerves in 3-month-old WT and *Stmn2* KO animals with quantification of nerve density (#/µm^2^) shown to the right. Scale bar= 25 µm. Statistical significance determined by Student’s unpaired t-test. **B)** Intraepidermal nerve fiber density of footpad skin of 3-month-old WT and *Stmn2* KO mice. Nerve fiber density quantified as # of nerves per mm of basement membrane. Statistical significance determined by Student’s unpaired t-test. **C)** Representative images from lumbrical muscles of 3-month-old WT and *Stmn2* KO mice. NMJs visualized by neurofilament/SV2 (green) and bungarotoxin (magenta). White arrow shows fragmented AChR clusters, periwinkle arrow indicates sprout from an adjacent NMJ. Quantification is based on the innervation of a region of AChR clusters contained within a 20X field. Scale bar = 25 µm. (WT animals n =7, KO n=10) **D)** Representative images from soleus and extensor digitalis longus (EDL) muscles from 3-month-old WT (n=3-4) and *Stmn2* KO (n=4) mice. Scale bar = 10 µm. Quantification of percent NMJs of each innervation status on the right. Arrow highlights denervated endplate. **E)** Representative electron micrographs of NMJs from 3-month-old WT and *Stmn2* KO lumbricals. Scale bar = 1 µm. Black arrow = junctional folds, blue arrow = presynaptic mitochondria, yellow arrow = synaptic vesicles (or lack thereof), * = postsynaptic mitochondrial clustering, SC = Schwann cell, N = myocyte nuclei, AR = axon remnants **F)** Quantification of synaptic vesicle density in the presynapse (#/µm^2^). Statistical significance determined by unpaired student’s test. Mitochondrial Circularity quantified on the right. “1” represents a perfect circle. Unless otherwise stated, statistical significance was determined using 2-way ANOVA with Sidak’s multiple comparisons test. (ns: not significant, *p<0.05, **p<0.01, ***p<0.001, ****p<0.0001).

The lack of axonal loss coupled with the robust behavioral abnormalities in *Stmn2* KO mice prompted us to examine the most distal units of both the motor and sensory systems. We first analyzed the intraepidermal nerve fiber (IENF) density in the footpad using PGP9.5 immunostaining and counted the fibers that crossed the basement membrane (Ebenezer *et al*., 2007). In *Stmn2* KO mice, IENF density (number per millimeter of dermis) in the mouse footpad is significantly reduced (Fig. 3B). Thus, we can conclude that STMN2 loss results in a distal sensory neuropathy affecting small unmyelinated nerve fibers in the footpad skin.

With predominantly motor deficits in ALS, we were most interested in evaluating the motor synapses, which have been implicated in early ALS pathology (Fischer *et al*., 2004; Dadon-Nachum *et al*., 2010). We examined NMJs from lumbrical muscles in the hind feet. In samples from wild type mice, we observe clustered bungarotoxin-labelled acetylcholine receptors forming characteristic postsynaptic endplates that are consistently co-localized with SV2-labelled presynapses. Strikingly, in the *Stmn2* KO mouse, the NMJ is severely disorganized with little innervation of AChRs and substantial endplate fragmentation. The individual AChR clusters were so aberrant that we could not quantify percent innervation because it was not possible to determine the relationship between the residual AChR clusters and presynaptic axons/terminals. We uniformly observe that AChR clusters appear fragmented, with many small clusters that do not form the characteristic pretzel morphology of wild type AChR clusters. When assessing fields of these AChR clusters, we would either observe A) a complete absence of any presynaptic elements, B) occasional motor axons in the vicinity of some AChR clusters, or C) apparently innervated endplate fragments with a dramatic presynaptic sprouting phenotype. Such sprouting is diagnostic of denervation of the endplate and subsequent reinnervation by regenerating fibers from adjacent NMJs (Brown &Ironton, 1978). Hence, the endplate fragmentation and presynaptic sprouting observed in *Stmn2* KO animals is consistent with presynaptic degeneration (Fig. 3C).

In ALS patients, there is preferential degeneration of fast-fatigable (FF) motor units (Dengler *et al*., 1990). Lumbrical muscles are primarily comprised of FF motor units; thus, we investigated whether other muscles primarily innervated by FF motoneurons are also susceptible to denervation in *Stmn2* KO mice. The extensor digitalis longus (EDL) is a fast twitch muscle located on the lateral side of the lower leg. The soleus muscle, which primarily consists of slow twitch muscle fibers, is located on the posterior portion of the lower leg at roughly the same distal position as the EDL and is relatively spared in many ALS mouse models. Neither of these muscles showed endplate fragmentation so we were able to quantify individual NMJs. For blinded quantification, we defined fully innervated NMJs as those with complete apposition of AChRs and synaptic vesicles, partially innervated as those with some presynapse apposition but with some exposed AChRs as well, and un-innervated as those with endplates but no presynapse present. Examining these muscles, we found denervation at the NMJs of the *Stmn2* KO EDL muscle, with more than half of all postsynaptic AChR clusters un-innervated. In contrast, denervation is not observed in the soleus muscle (Fig. 3D). While there is dramatic denervation of the EDL, it is less severe than the endplate fragmentation present in the more distal lumbrical muscles. Taken together, we conclude that the absence of STMN2 results in a motor neuropathy with NMJ defects that preferentially affect distal, fast-fatigable motor units.

With such a striking phenotype in lumbrical muscles, we next performed transmission electron microscopy (TEM) to explore the ultrastructural basis of the NMJs defects in three-month old *Stmn2* KO animals. In WT animals, NMJs are abundant, with clear apposition of presynaptic terminals opposite postsynaptic endings with characteristic junctional folds. These wildtype presynaptic terminals are densely occupied by synaptic vesicles and mitochondria, the postsynapse exhibits mitochondrial clustering near the edge of the myocyte and synapse, and Schwann cells are present nearby (Fig. 3E). The micrographs of NMJs from *Stmn2* KO mice are consistent with the observations from the immunofluorescence images— it is very difficult to identify NMJs on these muscles. Occasionally putative remnants of postsynaptic specializations are observed, with myocyte nuclei at the edge of muscle fibers and a cluster of mitochondria nearby, however there is rarely a recognizable presynaptic terminal. Instead, there are only remnants of an axon terminal with an occasional Schwann cell present. Those few NMJs that are still intact exhibit more circular mitochondria and synaptic vesicle depletion in the presynaptic terminal. In these cases, Schwann cells are present near the NMJ and the postsynapse appears more organized with some mitochondrial clustering at the edge of the myocyte (Fig. 3E,F). These ultrastructural findings are consistent with the behavioral, electrophysiological, and gross anatomical defects described above, all highlighting that STMN2 is required for maintenance of distal motor synapses.

### Stmn2 KO mouse motor phenotype is not due to SARM1 activation

We previously demonstrated that STMN2 is an axonal maintenance factor whose overexpression delays axon degeneration following injury (Shin *et al*., 2012). Furthermore, STMN2 co-localizes with the potent axonal maintenance factor NMNAT2 in transport vesicles, and is co-regulated with NMNAT2 by MAPK stress signaling and when overexpressed with NMNAT2 leads to synergistic axon protection (Gilley *et al*., 2010; Summers *et al*., 2018; Summers *et al*., 2020; Walker *et al*., 2017). Importantly, NMNAT2 inhibits SARM1, the central executioner of axon degeneration (reviewed in Figley &DiAntonio, 2020). We therefore hypothesized that STMN2 regulates NMNAT2 levels and that STMN2 loss causes the observed neurodegenerative phenotypes by reducing axonal NMNAT2 and promoting SARM1 activation.

To obtain large enough cell numbers for the biochemical and metabolic analysis necessary to test our hypothesis, we used CRISPR to knockdown *Stmn2* in cultured DRG neurons. We developed highly effective guide RNAs against *Stmn2* and used lentiviruses to express them in neurons from Cas9 transgenic mice. Contrary to our hypothesis, CRISPR/Cas9 KD of *Stmn2* in cultured neurons does not change axonal NMNAT2 levels as measured by Western blot (Fig. 4A,B). Levels of cADPR, a specific biomarker of SARM1 activity (Sasaki *et al*., 2020), are not significantly different in axonal lysates in neurons expressing STMN2 vs. scrambled control gRNAs (Fig. 4C), demonstrating that loss of STMN2 does not activate SARM1 *in vitro*.

**Figure 4:**
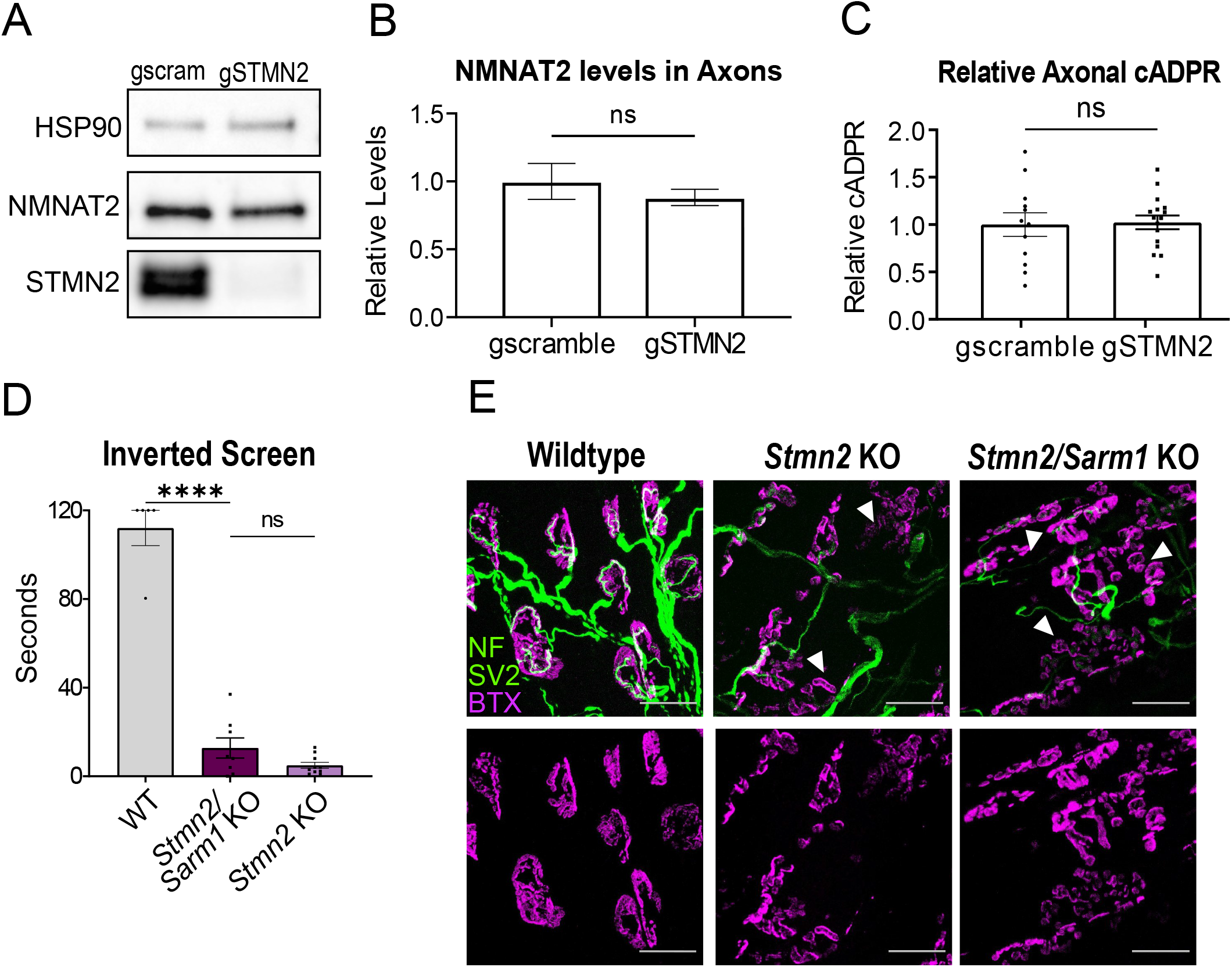
Phenotypes associated with STMN2 loss are not mediated by SARM1 activation. **A)** Representative Western blots of NMNAT2 and STMN2 using axonal lysates from neurons derived from Cas9 transgenic mice infected with lentivirus expressing *Stmn2* or scrambled control gRNAs. **B)** Relative NMNAT2 protein band intensity (n=4 experiments). **C)** Average relative cADPR levels from axonal lysates from neurons derived from Cas9 transgenic mice infected with lentivirus expressing *Stmn2* or scrambled control gRNAs. Statistical significance determined by Student’s unpaired t-test. **D)** Latency time to fall from an inverted screen (max. 120s) for wildtype, *Stmn2/Sarm1* KO, and *Stmn2* KO (from fig 2.) animals. Statistical significance determined by one-way ANOVA with Tukey’s multiple comparison test. **E)** Representative images of lumbrical NMJs from wildtype, *Stmn2/Sarm1* KO, and *Stmn2* KO mice. NMJs visualized by neurofilament/SV2 (green) and bungarotoxin (magenta). Arrows point to fragmented and denervated AChR clusters. Scale bar = 25 µm. (ns: not significant, ****p<0.0001)

To test the contribution of SARM1 to STMN2 phenotypes *in vivo*, we generated mice doubly mutant for *Sarm1* and *Stmn2* KO and assessed their motor phenotypes. At three months of age, *Stmn2/Sarm1* KO animals display the same severe motor function impairment as *Stmn2* KO mice (Fig. 4D). Moreover, presynaptic and postsynaptic elements of lumbrical NMJs from *Stmn2/Sarm1* KO animals were indistinguishable from those of *Stmn2* KO mice, including endplate fragmentation and many regions of partial innervation (Fig. 4E). Therefore, we conclude that SARM1 is not required for the motor phenotype of *Stmn2* KO mice. Moreover, these data suggest that STMN2 does not regulate the activity of SARM1, but instead promotes axonal maintenance via a parallel pathway.

### NMJ Denervation and Fragmentation is due to STMN2 loss in motor neurons

In ALS patients, TDP-43 dysfunction leads to loss of STMN2 protein in motor neurons, and STMN2 constitutive KO mice have motor defects and NMJ degeneration. This suggests that STMN2 is required in the motor neuron. To test the role of STMN2 in motor neurons, we selectively deleted *Stmn2* by crossing a floxed *Stmn2* allele (see Methods and Fig. 5A) with ChAT-Cre, which expresses Cre-recombinase in motor neurons. Excision is highly efficient, as staining in the spinal cord reveals a dramatic reduction of STMN2 in motor neurons of the Cre expressing mice (Supplementary Fig. 4A). To determine if STMN2 loss in motor neurons induces motor neuropathy, we tested the mice on the inverted screen. Indeed, the mice positive for ChAT-cre showed significant defects in hangtime (Fig. 5B). We next examined the effects of motor neuron-specific STMN2 loss on the structure of the NMJ. We stained lumbrical muscles for presynaptic and postsynaptic elements in ChAT-Cre^+^/*Stmn2*^*F/F*^ mice and found that the NMJs closely resemble those in *Stmn2* KO mice where most lumbrical muscle fibers are partially denervated with highly fragmented postsynaptic endplates (Fig. 5C). Hence, STMN2 deletion in motor neurons recapitulates the NMJ phenotype found in constitutive STMN2 KO mice. These data indicate that there is a cell-autonomous requirement for STMN2 in motoneurons, and that loss of STMN2 in motoneurons is sufficient to cause a distal motor neuropathy.

**Figure 5:**
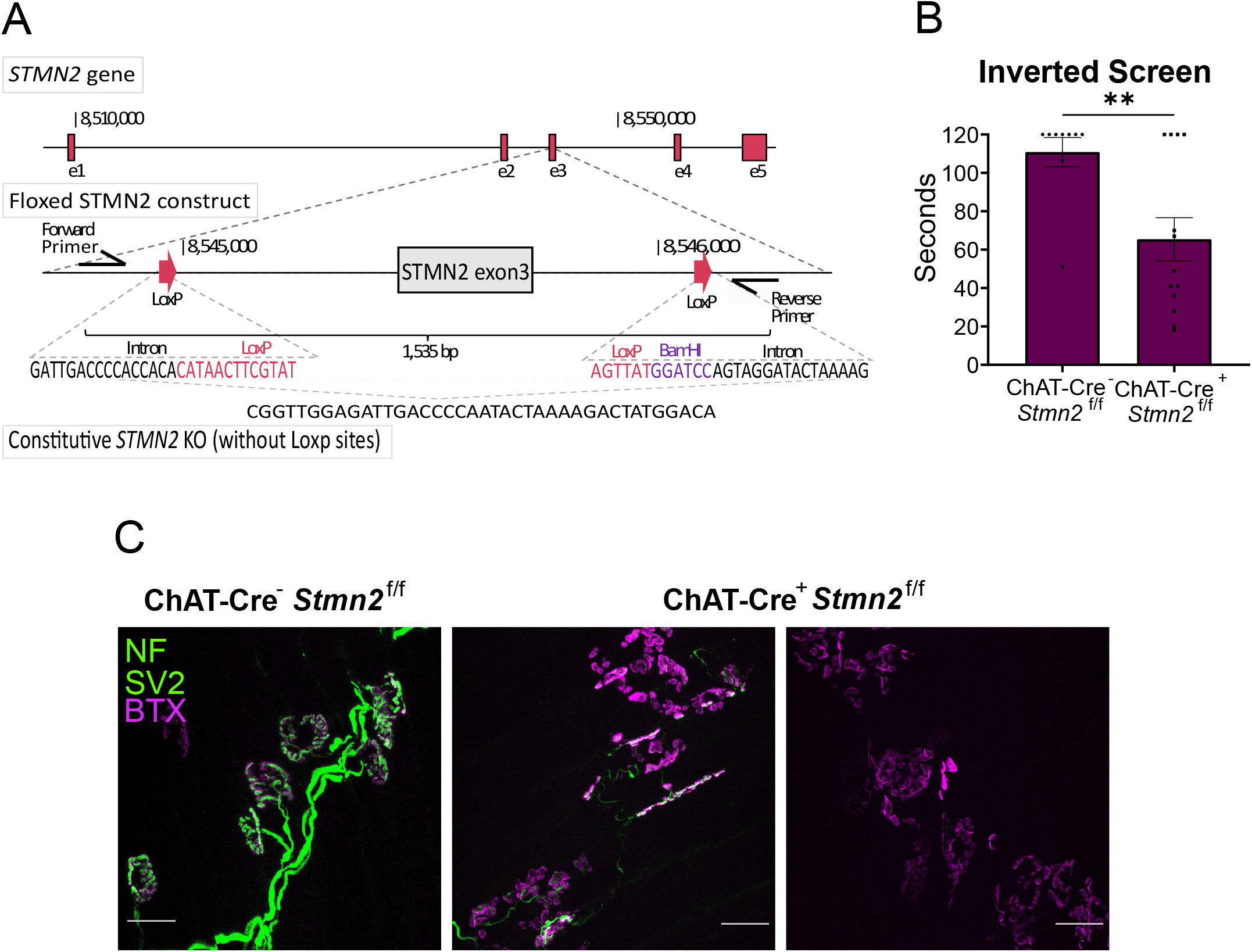
*Stmn2* KO in motor neurons causes motor pathology. **A)** Schematic of floxed *Stmn2* allele. **B)** Latency time to fall from an inverted screen (max. 120s) for ChAT-Cre^-^/*Stmn2*^*f/f*^ and ChAT-Cre^+^/*Stmn2*^*f/f*^ mice at 3 months of age. Statistical significance was determined by Student’s unpaired t-test. **C)** Representative images of NMJs from 3-month-old ChAT-Cre^-^/*Stmn2*^f/f^ and ChAT-Cre^+^/*Stmn2*^f/f^ lumbrical muscles. NMJs were visualized by neurofilament/SV2 (green) and bungarotoxin (magenta) staining. Scale bar = 25 µm. (**p<0.01)

### Stmn2^+/-^ heterozygous mice exhibit a progressive motor neuropathy

In both human neurons and patients with TDP-43 dysfunction, there is a reduction rather than complete absence of STMN2 (Klim *et al*., 2019; Melamed *et al*., 2019). Thus, to better model the partial loss of STMN2 that occurs in most cases of ALS, we studied the *Stmn2*^+/-^ heterozygous mice that show ∼50% reduction in STMN2 protein (Fig 1B). *Stmn2*^*+/-*^ heterozygous mice display normal motor function as young adults but develop a slowly progressive motor weakness by one-year of age (Fig. 6A). In contrast, one-year old *Stmn2*^*+/-*^ heterozygous mice behave normally in the Von Frey test (Fig. 6B) and rotarod (Fig. 6C). Hence, the partial loss of STMN2 protein results in a progressive and motor-selective behavioral defect.

**Figure 6:**
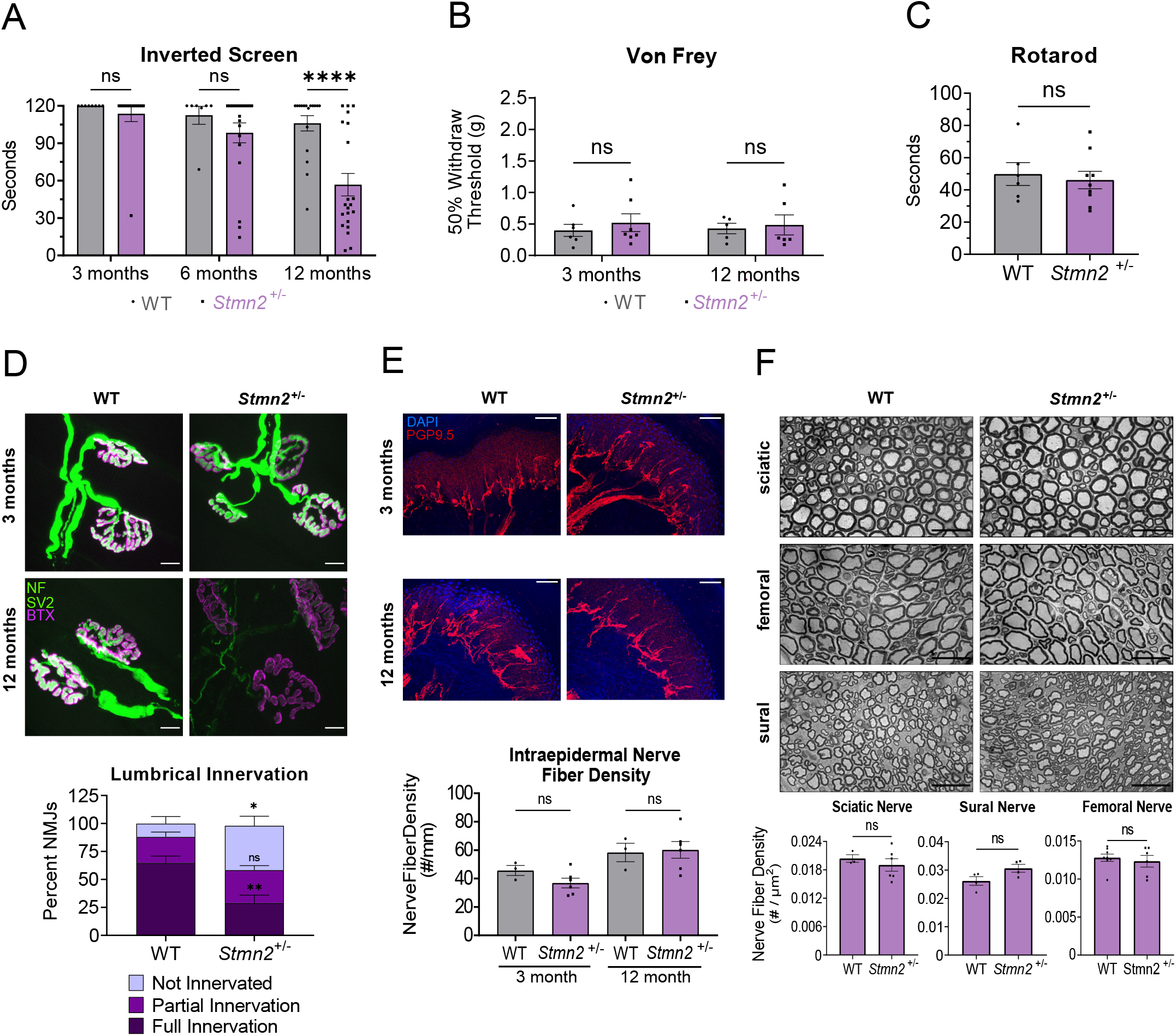
Partial STMN2 depletion results in progressive, distal motor neuropathy. **A)** Latency time to fall from an inverted screen (max. 120s) for WT and *Stmn2*^+/-^ mice at 3, 6, and 12 months of age. **B)** Average 50% hindpaw withdrawal threshold when force (grams) applied to 3 and 12-month-old WT and *Stmn2*^+/-^ mice. **C)** Time to fall (seconds) from an accelerating rod for 12-month-old WT and *Stmn2*^+/-^ mice. Statistical significance was determined by Student’s unpaired t-test. **D)** Lumbrical NMJs visualized by neurofilament/SV2 (green) and bungarotoxin (magenta) staining. Scale bar = 10 µm. Quantification is below. **E)** IENF labelled with anti-PGP9.5 and DAPI. Scale bar = 50µm. **F)** Representative images of sciatic, femoral, and sural nerves in 12-month-old WT and *Stmn2*^+/-^ animals with quantification of nerve density (#/µm^2^) shown underneath. Scale bar = 25 µm. Statistical significance determined by Student’s unpaired t-test. Unless otherwise noted, statistical significance was determined by 2-way ANOVA with Sidak’s multiple comparisons test. (ns: not significant, *p<0.05, **p<0.01, ****p<0.0001)

We next investigated the pathological basis for the motor dysfunction in one-year old *Stmn2*^*+/-*^ heterozygous mice. We focused on innervation of the lumbrical muscles, which were the most dramatically affected in *Stmn2* KO mice. Consistent with the absence of a motor phenotype at three months of age, staining *Stmn2*^*+/-*^ heterozygous and littermate control lumbrical muscle NMJs demonstrates postsynaptic AChR clusters with the classic pretzel morphology and well-apposed presynaptic terminals. Hence, NMJ development and maintenance appear normal in young adults. At one-year of age, postsynaptic AChR clustering still appears very similar to wild type at the lumbrical muscle NMJs of *Stmn2*^*+/-*^ heterozygous mice. Excitingly, however, the one-year-old *Stmn2*^*+/-*^ heterozygous mice exhibit denervation of lumbrical NMJs with a nearly four-fold increase in the fraction of fully denervated postsynaptic AChR clusters (Fig. 6D). Denervation occurs in roughly contiguous regions that are adjacent to well-innervated portions of the same muscle (Supplementary Fig. 5A), likely reflecting the degeneration of an axon branch. This phenotype is less severe than in the full *Stmn2* KO as the structure of the postsynapse is maintained, likely reflecting the ability of lower levels of STMN2 protein to provide some synaptic maintenance function. Consistent with this finding, we observe no obvious synaptic denervation in the more proximal EDL muscle of 12-month-old mice (Supplementary Fig. 5B), a muscle that also shows less dramatic denervation in the constitutive *Stmn2* KO. Constitutive *Stmn2* KO leads to loss of IENFs, so we tested whether this phenotype was also induced with partial loss of STMN2. We examined footpads of control and *Stmn2*^*+/-*^ mice and observed no difference in IENF density, demonstrating that motor endings are selectively vulnerable to reduction in STMN2 levels. (Fig. 6E). Finally, we observe no difference in the number of axons in the sural, sciatic or femoral nerves of *Stmn2*^*+/-*^ heterozygous mice, consistent with findings in the full *Stmn2* KO mice (Fig. 6F). Taken together, these findings demonstrate that a reduction in STMN2 protein levels leads to a progressive, distal-predominant motor neuropathy with NMJ denervation. These data strongly support the hypothesis that reduction in STMN2 protein levels contribute to ALS pathology and suggest that restoration of normal STMN2 protein levels in ALS patients with TDP-43 pathology could promote synaptic maintenance and motor function.

## Discussion

TDP-43 pathology occurs in almost all ALS patients, inspiring efforts to understand the consequences of TDP-43 dysfunction (Prasad *et al*., 2019). Two recent landmark studies identified *STMN2* as the most dysregulated transcript in TDP-43 deficient human motor neurons. They demonstrated that TDP-43 represses the inclusion of a cryptic exon in *STMN2* mRNA and found that STMN2 protein levels are reduced in most ALS patient spinal cords. These findings lead to the compelling hypothesis that loss of STMN2 is an important contributor to ALS pathology (Melamed *et al*., 2019; Klim *et al*., 2019). Here we test this hypothesis by generating and analyzing constitutive and conditional *Stmn2* knockout mice. We find that constitutive STMN2 loss induces an early motor and sensory neuropathy with distal NMJ denervation and concomitant motor defects. Fast-fatigable motor synapses, which are preferentially susceptible in ALS (Dengler *et al*., 1990), are also preferentially lost in these mice. STMN2 is required in motor neurons for synaptic maintenance, as selective excision of *Stmn2* in ChAT-expressing neurons triggers motor neuropathy and NMJ degeneration. To better model the partial loss of STMN2 protein that occurs in ALS, we examined *Stmn2* heterozygous mice. Excitingly, this moderate reduction in *Stmn2* expression is sufficient to cause a slowly progressive, motor-selective neuropathy with NMJ denervation. These findings strongly support the hypothesis that decreased STMN2 levels contribute to ALS pathology and raise the hope that increasing the levels of STMN2 protein in motor neurons will provide therapeutic benefit for ALS patients.

### *STMN2 Loss* in vivo *Recapitulates Aspects of ALS Neuromuscular Pathology*

ALS is a complex neurodegenerative disease with progressive loss of both upper and lower motor neurons. Symptoms often begin with distal motor weakness and neuromuscular degeneration, with gradual progression to more proximal muscles (van Es *et al*., 2017). The *Stmn2* KO mouse displays some aspects of this ALS pathology. First, STMN2 loss leads to distal weakness, as these mice have almost no capacity to hang on to the inverted screen, and yet have grossly normal gait reflecting the function of more proximal muscles. Second, *Stmn2* KO mice have distal-predominant NMJ loss. In the hind paw lumbrical muscles, we observe dramatic denervation with AChR cluster fragmentation. The more proximal EDL muscle shows less severe denervation, with denervated postsynaptic endplates retaining their classic pretzel-shaped morphology. Third, the *Stmn2* KO mice show selective loss of the fast-fatigable motor units which generally innervate fast twitch muscles. The EDL, a predominantly fast twitch muscle, displays extensive denervation, while the slow-twitch soleus muscle shows no denervation. The selective vulnerability of fast-fatigable motor units is seen both in patients with ALS (Dengler *et al*., 1990) and in mouse ALS models carrying pathogenic mutations in *FUS* (Sharma &Lyashchenko *et al*., 2016; Korobeynikov *et al*., 2022), SOD1 (Frey *et al*., 2000), and TDP-43 (Spiller *et al*., 2016, Ebstein *et al*., 2019). Finally, the *Stmn2* KO mice display sensory defects. While ALS is, of course, a predominantly motor disease, ALS patients can have sensory abnormalities as well. These symptoms can be accompanied by loss of intraepidermal nerve fibers (Riancho *et al*., 2021), consistent with the deficits we observe in *Stmn2* KO mice. While *Stmn2* deletion recapitulates important hallmarks of ALS, the mouse model does not exhibit axon loss or motor neuron death by one year of age.

Analysis of the *Stmn2* heterozygote revealed two additional aspects of the STMN2 phenotype that are consistent with ALS pathology. First, this more modest reduction in STMN2 protein results in a *progressive* motor neuropathy. As young adults, these mice perform normally on the inverted screen test, but by one-year of age show clear weakness. Similarly, young adult *Stmn2* heterozygous mice have fully innervated NMJs at their distal lumbrical muscles yet have lost many of these synapses by one-year. Second, *Stmn2* heterozygous mice reveal a motor selective neuropathy. In ALS, the motor system is preferentially affected. The *Stmn2* heterozygous mice show only motor deficits, with no loss of IENFs or sensory abnormalities. Taken together, the phenotypes of the heterozygous and homozygous *Stmn2* mutants are consistent, demonstrating that STMN2 is required for maintenance of distal NMJs, and imply that the dose of STMN2 is important, with the homozygote displaying an earlier and more severe disorder.

ALS is a highly heterogenous disease despite the nearly uniform presence of TDP-43 inclusions, indicating that there must be disease modifying factors. Thus, the dosage-sensitivity of the STMN2 loss-of-function phenotype has important implications for ALS pathogenesis and treatment. It implies that ALS disease progression may be influenced by the degree of STMN2 loss that results from TDP-43 dysfunction. More dramatic reductions in STMN2 could be associated with earlier age-of-onset and a more rapid progression of motor dysfunction and denervation. If so, then factors regulating STMN2 expression levels such as *STMN2* sequence variants (Theunissen *et al*., 2021) or natural variation in STMN2 transcription may be important modifiers of ALS disease course. We previously demonstrated that the neuronal stress kinase DLK (dual leucine zipper kinase) is a key regulator of STMN2 protein turnover, with activation of DLK leading to more rapid loss of axonal STMN2 (Summers *et al*., 2020). Interestingly, DLK is itself implicated in ALS pathogenesis, with activation of DLK stress signaling occurring in both ALS mouse models and, likely, in human patients (Le Pichon *et al*., 2017). If the DLK stress response were activated in TDP-43 dysfunctional neurons, it would lead to an even more dramatic loss of STMN2 from the axon. While there has been much interest in DLK as a therapeutic target for ALS, a recent clinical trial of a DLK inhibitor unfortunately failed due to its poor safety profile (Katz *et al*., 2022).

While DLK inhibition is likely not a viable approach for augmenting STMN2 function in ALS, the dosage sensitivity of the mouse phenotypes implies that boosting STMN2 levels could be therapeutic. One attractive approach is to use antisense oligonucleotides to block the cryptic splice site in neurons afflicted with dysfunctional TDP-43 (Klim *et al*., 2021). Alternatively, STMN2 gene therapy could be considered. In addition to boosting STMN2 levels, it is also worth exploring whether Stathmin-2 orthologs could provide therapeutic benefit. While we did not see ortholog upregulation in *Stmn2* KO mice, there still may be partial redundancy among the four stathmin paralogs, all of which are highly expressed in neurons (Ozon *et al*., 1998). Indeed, *Stmn1* mutants have a late-onset central and peripheral axonopathy, implying that STMN1 is also an axon maintenance factor (Liedtke *et al*., 2002). Furthermore, neuronal loss of the only *Drosophila* stathmin protein triggers NMJ degeneration (Graf *et al*., 2011), indicating that NMJ maintenance is a fundamental function of stathmin proteins rather than a unique attribute of STMN2. If so, then upregulation of any of the other STMNs could provide therapeutic benefit. Finally, TDP-43 pathology occurs in multiple neurodegenerative diseases. For example, TDP-43 inclusions have been seen in over half of individuals with Alzheimer Disease and presence of inclusions corresponds with greater cognitive impairment (Nag *et al*., 2018; Meneses *et al*., 2021). TDP-43 inclusions are also found in ∼50% of patients with frontotemporal dementia (Cairns *et al*., 2007). Importantly, the aberrant splice form of STMN2 is present in brains from FTD patients, (Prudencio *et al*., 2020) and STMN2 is decreased in AD brains (Mathys *et al*., 2019). Hence, methods for boosting stathmin function may also be beneficial in those devastating diseases.

STMN2 has been well described to regulate microtubule dynamics through binding of tubulin heterodimers to promote microtubule catastrophe (Reiderer *et al*., 1997). During development, this allows for growth cone guidance and axon outgrowth (Morii H *et al*.,2006). However, the mechanism by which STMN2 promotes synaptic maintenance after development is unknown. STMN2 is an axon maintenance factor in cultured neurons (Shin *et al*., 2012). Consistent with our current *in vivo* findings, this STMN2 loss was not sufficient to induce axon loss *in vitro*, however STMN2 did promote the stability of injured axons. Furthermore, STMN2 protein is co-regulated with the potent axonal maintenance factor and SARM1 inhibitor, NMNAT2 (Summers *et al*., 2018; Shin *et al*., 2012; Summers *et al*., 2020; Geisler *et al*., 2019; Walker *et al*., 2017). SARM1 is the central executioner of pathological axon degeneration (reviewed in Figley &DiAntonio, 2020), and rare activating mutations in SARM1 are enriched in ALS patients (Gilley *et al*., 2021; Bloom *et al*., 2022). Therefore, we hypothesized that STMN2 acts in concert with NMNAT2 to inhibit SARM1. However here we show that loss of STMN2 does not regulate NMNAT2 levels or promote SARM1 activation, and deletion of SARM1 does not suppress the STMN2 motor neuropathy phenotype in double-mutant mice. Hence, STMN2 must function in a parallel pathway to support axon/synapse survival. We previously demonstrated in both *Drosophila* and in cultured mammalian sensory neurons that stathmin is required for general axon transport and mitochondrial transport, respectively (Graf *et al*., 2011; Shin *et al*., 2012). Here we show that in *Stmn2* KO cultured neurons microtubule dynamics are disrupted, and electron micrographs revealed damaged mitochondria such as is seen in ALS (Atsumi, 1981). Mutations associated with ALS disrupt axonal transport (Castellanos-Montiel *et al*., 2020), and we suggest that this disruption of microtubule-based fast axonal transport may be the mechanism by which STMN2 deficiency disrupts the NMJ. Dysfunctional mitochondria could result from axonal transport deficits and could further potentiate NMJ instability.

This study demonstrates that loss of STMN2 recapitulates important aspects of ALS pathology. The finding that STMN2 loss does not replicate all aspects of ALS pathology is expected, as TDP-43 regulates the splicing of many neuronal transcripts. Indeed, two exciting new studies demonstrated TDP-43 represses cryptic exon inclusion in *UNC13A* transcripts (Ma *et al*., 2022; Brown *et al*., 2022). UNC13A is an important synaptic protein, and a non-coding SNP in *UNC13A* that affects splicing efficiency is significantly enriched among individuals with sporadic ALS (van Es *et al*., 2009, Ling *et al*., 2015). Likely, the combined depletion of STMN2, UNC13A, and other TDP-43 targets would result in a more complete model of ALS-like phenotypes. Moreover, TDP-43 has functions beyond regulating splicing, and its pathological inclusion in cytosolic aggregates may cause additional gain-of-function phenotypes (Prasad *et al*., 2019). While this model of TDP-43-dependent pathology suggests that combination treatments may be necessary to ameliorate ALS, the demonstration that STMN2 is a functionally relevant target is an important step forward for the development of rational treatments for this devastating disease.

## Materials and Methods

### Generation of *Stmn2* KO mice

All animal experiments were performed under the direction of institutional animal study guidelines at the Washington University, St. Louis, MO. The creation of the floxed *Stmn2* allele by the Genome Engineering &Stem Cell Center (GESC@MGI) via CRISPR (Clustered Regularly Interspaced Short Palindromic Repeats)/Cas9-mediated genetic modification (Chen *et al* 2011) was previously described (Sentmanat *et al*, 2022). In brief, guide RNAs (gRNAs) were designed corresponding to the introns flanking exon 3 and single-stranded oligodeoxynucleotides were designed with a loxP site and a BamHI site inserted directly in the gRNA cleavage sites, flanked by 60 bases of homology on each side. Transgenic C57BL/6N founders were generated by single-cell embryo electroporation. The constitutive KO allele was generated spontaneously in this process and confirmed by next generation sequencing.

### Mouse dorsal root ganglia neuron cultures

Embryonic DRG spot cultures were prepared as described by Gerdts *et al*. (Gerdts *et al*., 2013). Briefly, DRG neurons were collected from Cas9 knock-in mice (Jackson Laboratory) or WT and *Stmn2* KO littermates on embryonic day 12.5-13.5, genotyped, stored in Hibernate-A Medium (Thermo Fischer) overnight, and cultured the following day on plates coated with poly-D-lysine and laminin in neurobasal medium (Invitrogen) supplemented with 2% B27 (Invitrogen), 50 ng/mL nerve growth factor (Harlan Laboratories), and 1 uM 5-fluoro-2’-deoxyuridine (Sigma). Polymerization tracking studies were conducted on DIV6-7. Axon outgrowth assays were conducted on DIV3. Cell lysates were collected for Western blot and mass spectrometry on DIV 7.

### Lentivirus Transduction &Plasmids

EB3-mNeonGreen was cloned into the FCIV plasmid. EB3-mNeonGreen was a gift from Dorus Gadella (Addgene plasmid # 98881). Guide RNAs were cloned into a lentiguide vector. LentiGuide-Puro was a gift from Feng Zhang (Addgene plasmid # 52963). Sequences for STMN2 guide RNAs are: AGGTGAAGCAGATCAACAAC and GAAGAAAGACCTGTCTCTGG. The sequence for the scramble guide is: CGCGGCAGCCGGTAGCTATG. Lentiviruses were generated as previously described (Araki *et al*., 2004). Virus was added to Cas9 DRG cultures on DIV3. Virus was added to WT and *Stmn2* KO cultures on DIV1-2, achieving ∼100% transduction efficiency in DRG neurons.

### Cellular Imaging

#### Live Cell Microtubule Polymerization

Spot cultures were plated on FluoroDishes (World Precision Instruments, FD35-100) and transduced with the EB3-mNeonGreen lentivirus on DIV1-2. Live cell images were taken on a Nikon Spinning Disk Confocal Microscope every 3 seconds for 5 minutes. Kymographs were produced with Kymolyzer (Basu H *et al*., 2020) and analyzed with KymoButler (Jakobs MAH *et al*., 2019).

#### Axon Outgrowth

Spot cultures were grown in 4 well chamber glass slides so that WT and KO cultures were in adjacent wells. On DIV 3, cells were fixed for 15 minutes with 4% PFA. The cells were washed with PBS, blocked in 10% NGS 0.1% triton X-100 in PBS (PBS-T), and incubated with anti-TUJ1 primary antibody diluted in blocking buffer overnight at 4°C. The following day they were washed with PBS-T and incubated with Alexa Fluoro 488 conjugated goat anti-rabbit secondary (1:1000, Invitrogen, A11034) for one hour at room temperature. Afterwards, they were washed and mounted with Vectashield mounting media. Images were taken on a Leica DMI 4000B Confocal Microscope. The distance between the perimeter of each cell spot and the outer perimeter of its axon halo were measured in four different locations in each image and averaged. *Stmn2* KO axon lengths are normalized to WT axon length in the same experiment.

### Western Blot

Cell and brain lysates were resolved using SDS polyacrylamide gel electrophoresis (PAGE) on 4-20% Mini-Protean Gel (BioRad), followed by immunoblotting for NMNAT2 (1:100 anti-NMNAT2 (B-10) mouse monoclonal IgG1, sc-515206), STMN1 (1:1000 Anti-Stathmin 1 Rabbit Monoclonal Antibody, ab52630), STMN2 (1:1000 anti-SCG10, Shin *et al*., 2012), STMN3 (1:2000 Anti-STMN3 Rabbit Polyclonal Antibody, 11311-1-AP), STMN4 (1:1000 Anti-STMN4 Rabbit Polyclonal, 12027-1-AP), HSP90 (1:1000 Anti-HSP90 Rabbit IgG, C45G5), and TUJ1(anti-B3 tubulin, 1:10000 Simga-Aldrich, T2200), and visualized using standard chemiluminescence. Band intensity was quantified using ImageJ. Blots were normalized to their respective loading controls and then normalized to the control sample.

### Mass Spectrometry

Sciatic nerves were homogenized in 160 uL of cold 50% MeOH in water using a sonicator and then centrifuged (15000xg, 4°C, 10 minutes). The supernatant was transferred to a new tube with 50 µL choloroform. The mixture was shaken vigorously and centrifuged again (15000xg, 4°C, 10 minutes). This chloroform extraction was repeated twice. The clear aqueous phase was transferred to a new tube, lyophilized, and stored at -80°C until measurement. Lyophilized samples were reconstituted with 5mM ammonium formate (50µL, Sigma) and centrifuged for 10 minutes at 12000xg. Supernatant was transferred to the sample plate, and metabolite measurements were acquired as previously described (Sasaki *et al*., 2016). Axonal metabolites were collected on DIV7 by removing the cell body spot from the well and extracting metabolites as previously described (Sasaki *et al*., 2020).

### Behavioral Assays

#### Von Frey Assay

Von Frey assay was performed according to the up-down method (Chaplan *et al*., 1994). Briefly, mice were habituated individually in plexiglass boxes on a wire mesh screen. They were given four hours to rest before testing. The plantar surface of their hind paw was stimulated for 2 seconds with a filament, beginning at a bending force of 0.32 g. If withdraw occurred, a filament that delivered less force would be used next. If withdraw did not occur, a filament of greater force would be used. Stimulation occurred four more times after the first change in response. The 50% withdraw threshold was calculated (Dixon, 1980), and results from left and right hind paws were averaged for each mouse. Mice were allowed to rest between trials, and mice were always tested at the same time of day.

#### Inverted Screen Assay

Mice were place on a wire mesh screen, and the screen was inverted. Each mouse underwent 3 trials with five-minute rest periods in between, and the latency to fall for each mouse was recorded. If the mouse was still hanging on the screen at 120 seconds, they were taken off the screen and 120 seconds was recorded. The average of the three trials was taken.

#### Rotarod Assay

The mice were trained to walk on a rotarod (Panlab, LE8205) at a constant speed for five minutes the day prior to testing. On the day of testing, they were trained for another five minutes. Each mouse then underwent five trials on the rotarod that started at 4 rpm and accelerated every second with five minutes of rest in between. Latency to fall was recorded and the average of the five trials was taken.

### Nerve Electrophysiology

Compound muscle action potentials (CMAPs) were acquired using a Viking Quest electromyography device (Nicolet) as previously described (Beirowski *et al*., 2011). Briefly, mice were anesthetized, then a stimulating electrode was placed in the sciatic notch and a recording electrode was placed in the foot. Supramaximal stimulation was used for CMAPs. SNAPs were acquired using the same device as previously described (Geisler S *et al*., 2016). Mice were anesthetized and electrodes were placed subcutaneously with the stimulating electrode placed in the tail tip 30 mm distal from the recording electrode that was placed into the base of the tail. A ground electrode was placed in between the two. Supramaximal stimulation was used for SNAPs.

### Spinal Cord

Spinal cords were dissected and embedded in O.C.T., twenty-five-micron thick sections cut on a cryostat and mounted onto slides. Slides were dried at room temperature overnight, incubated in 0.1% Triton-X in PBS for 30 minutes, blocked in 4% BSA 1% Triton-X in PBS for 30 minutes, and incubated overnight at 4°C with goat anti-ChAT (1:100, Millipore Sigma-Aldrich, AB144P) and anti-STMN2 (1:100, rabbit anti-SCG10, Shin *et al*., 2012) in blocking buffer. The next day, slides were washed with 0.1% Triton-X 3 times, incubated with Cy3 anti-goat (1:250, Jackson Immunoresearch, 705-166-147) and Alexa Fluoro-488 goat anti-rabbit IgG (H+L) (Invitrogen, A11034) in wash solution for 2 hours at room temperature, then washed in PBS and mounted in Vectashield with DAPI. Images of motor neurons in the lumbar spinal cord were taken on a Zeiss Axio Imager Z2 Fluorescence Microscope with ApoTome 2. Cell bodies were counted and averaged across 3 or more sections. For quantification of STMN2, motor neurons were identified by ChAT expression and the percent of STMN2+ motor neurons was calculated.

### Nerve structural analysis

Sciatic, sural, and femoral nerves were processed as previously described (Geisler S *et al*., 2016). Briefly, nerves were fixed in 3% glutaraldehyde in 0.1 ml PBS overnight at 4°C, washed and stained with 1% osmium tetroxide (Sigma Aldrich) overnight at 4°C. Nerves were washed and dehydrated in a serial gradient of 50% to 100% ethanol. Nerves were then incubated in 50% propylene oxide/50% ethanol, then 100% propylene oxide. Following that, nerves were incubated in Araldite resin/propylene oxide solutions in 50:50, 70:30, 90:10 ratios overnight, and subsequently embedded in 100% Araldite resin solution (Araldite: DDSA: DMP30, 12:9:1, Electron Microscopy Sciences) and baked at 60°C overnight. Semithin 400—600 nm sections were cut using a Leica EM UC7 Ultramicrotome, placed on microscopy slides, and stained with Toluidine blue. Staining and quantification were performed as previously described. (Sasaki *et al*., 2018) All quantification was performed blinded.

### Intraepidermal nerve fiber density quantification

Intraepidermal nerve fiber (IENF) staining and quantification was performed as previously described (Geisler S *et al*., 2016). Briefly, the footpad skin was placed in fresh picric acid fixative overnight at 4°C. The following day the samples were washed in PBS and placed into 30% sucrose for over 24 hours. Samples were embedded in O.C.T. and sectioned using a cryostat in 50 micron sections and placed floating in cryoprotectant (30% sucrose and 33% ethylene glycol in PBS) and stored at -20°C. Free floating sections were then washed with PBS, blocked in 5% normal goat serum 0.3% Triton-X in PBS, then incubated in blocking solution with PGP9.5 antibody (1:1000, AB1761, Millipore) overnight at 4°C. The following day the free-floating sections were washed with PBS-T, incubated with anti-rabit-Cy3 (1:500) for 2 hours at room temperature, washed again with PBS-T, and mounted in Vectshield with DAPI. The footpads were imaged on a Leica DMI 4000B confocal microscope using a 20x objective. Z-stacks were acquired through the whole sample and maximal projection was applied. Density was quantified by the number of PGP9.5+ axons that crossed the basement membrane and normalized to the length of the basement membrane. Densities were average over 3 sections per animal. Imaging and analysis were performed blinded to genotype.

### NMJ Analysis

Dissected muscles were incubated in 2% Triton-X in PBS for 30 minutes, blocked with 5% BSA 1% Triton-X for 30 minutes, and incubated overnight at 4°C in blocking buffer with primary antibodies against neurofilament and SV2 (2H3, 1:100, DHSB AB2314897; SV2, 1:200, DSHB AB2315387). The following day, the samples were washed in PBS, incubated in 1% Triton-X in PBS with FITC rabbit anti-mouse IgG1 (1:400, Invitrogen A21121) and Alexa fluoro-568 conjugated a-bungarotoxin (1:500, Biotium 00006) for 3 hours at room temperature. Muscles were washed in PBS 3 times for 15 minutes and were mounted with Vectashield Mounting medium.

To analyze NMJ morphology, z-stack images were obtained using a confocal microscope and maximal projection was applied. Due to the severe disruption of endplate morphology observed in many lumbrical muscles, the innervation status of lumbrical muscles was determined as of images of areas containing postsynaptic endplates were obtained and each 20x field was categorized as 1) fully innervated, 2) partially innervated, 3) sprouting with endplate disruption, or 4) not innervated. Because other muscles had preserved endplate morphology, their innervation status was quantified by assessing individual NMJs. Individual NMJs were categorized as 1) fully innervated, 2) partially innervated, or 3) not innervated. Over 50 NMJs were evaluated per animal for soleus and EDL muscles. Images shown are representative images taken with a 63x oil immersion lens. Researchers were blinded to genotype during imaging and image analysis.

### Embryonic Tissue

Embryos were collected at embryonic day 18.5-19.5 and incubated decapitated in 4% PFA overnight. The following day, embryos were rinsed with PBS. Heads were placed in 30% sucrose for multiple days to ensure equilibration, then embedded in O.C.T. 18-micron sections were cut using a cryostat and collected on slides. Sections were stained according to the same protocol as adult spinal cords using anti-2H3 (1:300, DHSB AB2314897) and Alexa fluoro-568 anti-mouse (1:250, Thermo Scientific, A-21124). Images were taken using a 20x lens on a Zeiss Axio Scan Z1 slide scanner. Diaphragms were dissected from the body and stained free floating utilizing the protocol described above for adult muscles. Diaphragm images were taken using a 20x objective along the endplate band in the lateral, anterior portion of the diaphragm. ImageJ was used for quantification from two images per embryo.

### Transmission Electron Microscopy

Mice were perfused with 4% PFA and whole feet immersed in 3% glutaraldehyde for 3 days. Lumbrical muscles were then dissected out, rinsed with 0.1M phosphate buffer, placed in 1% osmium tetroxide overnight at 4°C, subsequently washed with 0.1M PB 3 times, and dehydrated. Dehydration consisted of 30-minute incubations with increasing concentrations of acetone in water: 50%, 70%, 90%, and finally three times in 100% acetone. Muscles were further incubated in Spurr:acetone prepared using the Spurr Resin kit (Electron Microscopy Science, 14300), at ratios of 1:2, 1:1, 3:1 and finally 100% Spurr, with each incubation lasting 24 hours at room temperature. Lumbricals were then embedded in 100% Spurr and polymerized at 65°C for 48 hours. Thin sections were cut by the Washington University Core for Cellular Imaging (WUCCI) and placed on screens for imaging. A JEOL JEM-1400Plus transmission electron microscope was used for imaging. 6-10 images were taken per sample. Presynapse mitochondrial circularity was quantified using the formula Circularity = 4π*area/perimeter^2^, as described (Joelcio *et al*., 2020). As the value approaches 0.0, it indicates an increasingly elongated polygon. A circularity value of 1.0 indicates a perfect circle. Additionally, synaptic vesicles in the presynapse were counted, and the number of vesicles was divided by the area to get synaptic vesicle density. Only mitochondria and synaptic vesicles from intact axons were quantified. The researcher was blinded to genotype during image analysis.

### Statistical Analysis

Data are reported as means ± standard error of the mean (SEM). Statistics were calculated with the aid of Prism 9 (Graphpad) software. For between group comparisons, one-way and two-way ANOVA were used with post-hoc Holm-Sidak multiple comparison or post-hoc Tukey’s multiple comparison tests and unpaired t-tests when appropriate. Two-tailed significance tests were used with p < 0.05 considered statistically significant.

## Acknowledgments

We would like to thank members of the DiAntonio and Milbrandt labs for their thoughtful discussions on the study. We would also like to thank Cassidy Menendez, Rachel McClarney, Alicia Neiner, Sylvia Johnson, Xiaolu Sun, Kelli Simburger, Matthew Figley, John Palucki, Yo Sasaki, and Liya Yuan for their technical support and/or technical training and advice. Thank you to Margaret Hayne for her guidance on writing and making figures. We also thank the Washington University Core for Cellular Imaging (WUCCI) for their technical support, expertise, and training on the spinning disk microscope and transmission electron microscope. We would also like to thank the Genome Engineering &Stem Cell Center (GESC@MGI) at Washington University for generating the *Stmn2* mice.

## Author Contributions

KLK, AJB, AD, and JM conceived the overall study. All authors contributed to the study design. KLK and AS collected mouse behavior data. KLK and AS collected mouse tissue samples and performed immunofluorescent staining on collected tissues. LD performed and analyzed live cell microscopy. YY and RES performed TEM and interpreted electron micrographs. YY analyzed electron micrographs. KLK performed all other cell culture experiments, metabolite and protein assays, confocal imaging, and image analysis. KLK wrote the manuscript and prepared all figures. AJB, AD, and JM oversaw the analysis and revised the manuscript. All authors gave final approval of the manuscript.

## Corresponding authors

Correspondence to JM and AD

**Supplement 1: Innervation of the diaphragm in developing *Stmn2* KO embryos**.

**A)** Representative images of diaphragm pre- and post-synaptic elements in WT and *Stmn2* KO E18.5 embryos, visualized by neurofilament/SV2 (green) and bungarotoxin (magenta) staining.

**B)** Quantification of neurites and AChR clusters. Statistical significance determined by Student’s unpaired t-test. (ns: not significant, ***p<0.001)

**Supplement 2: Stathmin orthologs are not upregulated in *Stmn2* KO brains** Representative Western blots and quantification (n=5-6 per genotype) of **A)** STMN1, **B)** STMN3, and **C)** STMN4 in *Stmn2* KO and *Stmn2*^+/-^ brains. TUJ1 and GAPDH were used as loading controls. (ns: not significant)

**Supplement 3: Intact Nervous System in *Stmn2* KO animals**

**A)** Representative images of sciatic, femoral, and sural nerves from 12 month old WT and *Stmn2* KO animals. **B)** Motor neurons in WT and *Stmn2* KO lumbar spinal cord sections are labelled with anti-ChAT antibodies. Quantification on the right. (ns: not significant)

**Supplement 4: Successful deletion of *Stmn2* in motor neurons**

**A)** Representative images of 3-month-old ChAT-Cre^-^/ *Stmn2*^*f/f*^ and ChAT-Cre^+^/*Stmn2*^*f/f*^ ventral horn motor neurons labelled with anti-ChAT (red) and anti-STMN2 (green) antibodies. The percent of STMN2+ motor neurons is quantified to the right. Statistical significance was determined using a Student’s unpaired t-test.

**Supplement 5: Innervation in *Stmn2***^**+/-**^ **mice**

**A)** Representative low-magnification image of *Stmn2*^+/-^ mouse NMJs on lumbrical muscles, highlighting the regional distribution of denervated areas from the *Stmn2*^+/-^ mouse. **B)** Representative images of WT and *Stmn2*^+/-^ mouse NMJs in EDL muscles with quantification below. Statistical significance was determined by 2-way ANOVA with Sidak’s multiple comparisons test. (ns: not significant)

## References

Anderson DJ, Axel R. Molecular probes for the development and plasticity of neural crest derivatives. Cell. 1985 Sep;42(2):649–62. doi: 10.1016/0092-8674(85)90122-9. PMID: 3839717.

Araki T, Sasaki Y, Milbrandt J. Increased nuclear NAD biosynthesis and SIRT1 activation prevent axonal degeneration. Science 2004;305: 1010–3.

Atsumi T. The ultrastructure of intramuscular nerves in amyotrophic lateral sclerosis. Acta Neuropathol. 1981;55(3):193–8. doi: 10.1007/BF00691318. PMID: 7349578.

Basu H, Ding L, Pekkurnaz G, Cronin M, Schwarz TL. Kymolyzer, a Semi-Autonomous Kymography Tool to Analyze Intracellular Motility. Curr Protoc Cell Biol. 2020 Jun;87(1):e107. doi: 10.1002/cpcb.107. PMID: 32530579.

Beirowski B, Gustin J, Armour SM, Yamamoto H, Viader A, North BJ, Michán S, Baloh RH, Golden JP, Schmidt RE, et al. Sir-two-homolog 2 (Sirt2) modulates peripheral myelination through polarity protein Par-3/atypical protein kinase C (aPKC) signaling. Proc Natl Acad Sci U S A. 2011 Oct 25;108(43):E952–61. doi: 10.1073/pnas.1104969108. Epub 2011 Sep 26. PMID: 21949390; PMCID: PMC3203793.

Bloom AJ, Mao X, Strickland A, Sasaki Y, Milbrandt J, DiAntonio A. Constitutively active SARM1 variants that induce neuropathy are enriched in ALS patients. Mol Neurodegener. 2022 Jan 6;17(1):1. doi: 10.1186/s13024-021-00511-x. PMID: 34991663; PMCID: PMC8739729.

Brown AL, Wilkins OG, Keuss MJ, Hill SE, Zanovello M, Lee WC, Bampton A, Lee FCY, Masino L, Qi YA, et al. TDP-43 loss and ALS-risk SNPs drive mis-splicing and depletion of UNC13A. Nature. 2022 Mar;603(7899):131–137. doi: 10.1038/s41586-022-04436-3. Epub 2022 Feb 23. PMID: 35197628.

Brown MC, Ironton R. Sprouting and regression of neuromuscular synapses in partially denervated mammalian muscles. J Physiol. 1978 May;278:325–48. doi: 10.1113/jphysiol.1978.sp012307. PMID: 671308; PMCID: PMC1282352.

Cairns NJ, Neumann M, Bigio EH, Holm IE, Troost D, Hatanpaa KJ, Foong C, White CL 3rd, Schneider JA, Kretzschmar HA, et al. TDP-43 in familial and sporadic frontotemporal lobar degeneration with ubiquitin inclusions. Am J Pathol. 2007 Jul;171(1):227–40. doi: 10.2353/ajpath.2007.070182. PMID: 17591968; PMCID: PMC1941578.

Castellanos-Montiel MJ, Chaineau M, Durcan TM. The Neglected Genes of ALS: Cytoskeletal Dynamics Impact Synaptic Degeneration in ALS. Front Cell Neurosci. 2020 Nov 13;14:594975. doi: 10.3389/fncel.2020.594975. PMID: 33281562; PMCID: PMC7691654.

Chaplan SR, Bach FW, Pogrel JW, Chung JM, Yaksh TL. Quantitative assessment of tactile allodynia in the rat paw. J Neurosci Methods 1994; 53: 55–63.

Chen F, Pruett-Miller SM, Huang Y, Gjoka M, Duda K, Taunton J, et al. High-frequency genome editing using ssDNA oligonucleotides with zinc-finger nucleases. Nat Methods. 2011;8(9):753–5.

Chertkova AO, Mastop M, Postma M, van Bommel N, van der Niet S, Batenburg KL, Joosen L, Gadella TWJ, Okada Y, Goedhart J. Robust and Bright Genetically Encoded Fluorescent Markers for Highlighting Structures and Compartments in Mammalian Cells. bioRxiv 160374 10.1101/160374

Dadon-Nachum M, Melamed E, Offen D. The “dying-back” phenomenon of motor neurons in ALS. J Mol Neurosci. 2011 Mar;43(3):470–7. doi: 10.1007/s12031-010-9467-1. Epub 2010 Nov 7. PMID: 21057983.

Dengler R, Konstanzer A, Küther G, Hesse S, Wolf W, Struppler A. Amyotrophic lateral sclerosis: macro-EMG and twitch forces of single motor units. Muscle Nerve. 1990 Jun;13(6):545–50. doi: 10.1002/mus.880130612. PMID: 2366827.

Di Paolo G, Lutjens R, Osen-Sand A, Sobel A, Catsicas S, Grenningloh G. Differential distribution of stathmin and SCG10 in developing neurons in culture. J Neurosci Res. 1997 Dec 15;50(6):1000–9. doi: 10.1002/(SICI)1097-4547(19971215)50:6<1000::AID-JNR10>3.0.CO;2-8. PMID: 9452014.

Dixon WJ. Efficient analysis of experimental observations. Annu Rev Pharmacol Toxicol 1980; 20: 441–62.

Ebenezer GJ, McArthur JC, Thomas D, Murinson B, Hauer P, Polydefkis M, Griffin JW. Denervation of skin in neuropathies: the sequence of axonal and Schwann cell changes in skin biopsies. Brain. 2007 Oct;130(Pt 10):2703–14. doi: 10.1093/brain/awm199. PMID: 17898011.

Ebstein SY, Yagudayeva I, Shneider NA. Mutant TDP-43 Causes Early-Stage Dose-Dependent Motor Neuron Degeneration in a TARDBP Knockin Mouse Model of ALS. Cell Rep. 2019 Jan 8;26(2):364-373.e4. doi: 10.1016/j.celrep.2018.12.045. PMID: 30625319.

Figley MD, DiAntonio A. The SARM1 axon degeneration pathway: control of the NAD^+^ metabolome regulates axon survival in health and disease. Curr Opin Neurobiol. 2020 Aug;63:59–66. doi: 10.1016/j.conb.2020.02.012. Epub 2020 Apr 17. PMID: 32311648; PMCID: PMC7483800.

Fischer LR, Culver DG, Tennant P, Davis AA, Wang M, Castellano-Sanchez A, Khan J, Polak MA, Glass JD. Amyotrophic lateral sclerosis is a distal axonopathy: evidence in mice and man. Exp Neurol. 2004 Feb;185(2):232–40. doi: 10.1016/j.expneurol.2003.10.004. PMID: 14736504.

Frey D, Schneider C, Xu L, Borg J, Spooren W, Caroni P. Early and selective loss of neuromuscular synapse subtypes with low sprouting competence in motoneuron diseases. J Neurosci. 2000 Apr 1;20(7):2534–42. doi: 10.1523/JNEUROSCI.20-07-02534.2000. PMID: 10729333; PMCID: PMC6772256.

Geisler S, Doan RA, Strickland A, Huang X, Milbrandt J, DiAntonio A. Prevention of vincristine-induced peripheral neuropathy by genetic deletion of SARM1 in mice. Brain. 2016 Dec;139(Pt 12):3092-3108. doi: 10.1093/brain/aww251. Epub 2016 Oct 25. PMID: 27797810; PMCID: PMC5840884.

Geisler S, Doan RA, Cheng GC, Cetinkaya-Fisgin A, Huang SX, Höke A, Milbrandt J, DiAntonio A. Vincristine and bortezomib use distinct upstream mechanisms to activate a common SARM1-dependent axon degeneration program. JCI Insight. 2019 Sep 5;4(17):e129920. doi: 10.1172/jci.insight.129920. PMID: 31484833; PMCID: PMC6777905.

Gerdts J, Summers DW, Sasaki Y, DiAntonio A, Milbrandt J. Sarm1-mediated axon degeneration requires both SAM and TIR interactions. J Neurosci. 2013 Aug 14;33(33):13569–80. doi: 10.1523/JNEUROSCI.1197-13.2013. PMID: 23946415; PMCID: PMC3742939.

Gilley J, Coleman MP. Endogenous Nmnat2 is an essential survival factor for maintenance of healthy axons. PLoS Biol. 2010 Jan 26;8(1):e1000300. doi: 10.1371/journal.pbio.1000300. PMID: 20126265; PMCID: PMC2811159.

Gilley J, Jackson O, Pipis M, Estiar MA, Al-Chalabi A, Danzi MC, van Eijk KR, Goutman SA, Harms MB, Houlden H, et al. Enrichment of SARM1 alleles encoding variants with constitutively hyperactive NADase in patients with ALS and other motor nerve disorders. Elife. 2021 Nov 19;10:e70905. doi: 10.7554/eLife.70905. PMID: 34796871; PMCID: PMC8735862.

Graf ER, Heerssen HM, Wright CM, Davis GW, DiAntonio A. Stathmin is required for stability of the Drosophila neuromuscular junction. J Neurosci. 2011 Oct 19;31(42):15026–34. doi: 10.1523/JNEUROSCI.2024-11.2011. PMID: 22016536; PMCID: PMC3207242.

Jakobs MA, Dimitracopoulos A, Franze K. KymoButler, a deep learning software for automated kymograph analysis. Elife. 2019 Aug 13;8:e42288. doi: 10.7554/eLife.42288. PMID: 31405451; PMCID: PMC6692109.

Katz JS, Rothstein JD, Cudkowicz ME, Genge A, Oskarsson B, Hains AB, Chen C, Galanter J, Burgess BL, Cho W, et al. A Phase 1 study of GDC-0134, a dual leucine zipper kinase inhibitor, in ALS. Ann Clin Transl Neurol. 2022 Jan;9(1):50–66. doi: 10.1002/acn3.51491. Epub 2022 Jan 10. PMID: 35014217; PMCID: PMC8791798.

Klim JR, Williams LA, Limone F, Guerra San Juan I, Davis-Dusenbery BN, Mordes DA, Burberry A, Steinbaugh MJ, Gamage KK, Kirchner R, et al. ALS-implicated protein TDP-43 sustains levels of STMN2, a mediator of motor neuron growth and repair. Nat Neurosci. 2019 Feb;22(2):167–179. doi: 10.1038/s41593-018-0300-4. Epub 2019 Jan 14. PMID: 30643292; PMCID: PMC7153761.

Klim JR, Pintacuda G, Nash LA, Guerra San Juan I, Eggan K. Connecting TDP-43 Pathology with Neuropathy. Trends Neurosci. 2021 Jun;44(6):424–440. doi: 10.1016/j.tins.2021.02.008. Epub 2021 Apr 5. PMID: 33832769.

Korobeynikov VA, Lyashchenko AK, Blanco-Redondo B, Jafar-Nejad P, Shneider NA. Antisense oligonucleotide silencing of FUS expression as a therapeutic approach in amyotrophic lateral sclerosis. Nat Med. 2022 Jan;28(1):104–116. doi: 10.1038/s41591-021-01615-z. Epub 2022 Jan 24. PMID: 35075293; PMCID: PMC8799464.

Le Pichon CE, Meilandt WJ, Dominguez S, Solanoy H, Lin H, Ngu H, Gogineni A, Sengupta Ghosh A, Jiang Z, Lee SH, et al. Loss of dual leucine zipper kinase signaling is protective in animal models of neurodegenerative disease. Sci Transl Med. 2017 Aug 16;9(403):eaag0394. doi: 10.1126/scitranslmed.aag0394. PMID: 28814543.

Liedtke W, Leman EE, Fyffe RE, Raine CS, Schubart UK. Stathmin-deficient mice develop an age-dependent axonopathy of the central and peripheral nervous systems. Am J Pathol. 2002 Feb;160(2):469–80. doi: 10.1016/S0002-9440(10)64866-3. PMID: 11839567; PMCID: PMC1850667.

Ling JP, Pletnikova O, Troncoso JC, Wong PC. TDP-43 repression of nonconserved cryptic exons is compromised in ALS-FTD. Science. 2015 Aug 7;349(6248):650–5. doi: 10.1126/science.aab0983. PMID: 26250685; PMCID: PMC4825810.

Ling SC, Polymenidou M, Cleveland DW. Converging mechanisms in ALS and FTD: disrupted RNA and protein homeostasis. Neuron. 2013 Aug 7;79(3):416–38. doi: 10.1016/j.neuron.2013.07.033. PMID: 23931993; PMCID: PMC4411085.

Ma XR, Prudencio M, Koike Y, Vatsavayai SC, Kim G, Harbinski F, Briner A, Rodriguez CM, Guo C, Akiyama T, et al. TDP-43 represses cryptic exon inclusion in the FTD-ALS gene UNC13A. Nature. 2022 Mar;603(7899):124–130. doi: 10.1038/s41586-022-04424-7. Epub 2022 Feb 23. PMID: 35197626.

Mathys H, Davila-Velderrain J, Peng Z, Gao F, Mohammadi S, Young JZ, Menon M, He L, Abdurrob F, Jiang X, et al. Single-cell transcriptomic analysis of Alzheimer’s disease. Nature. 2019 Jun;570(7761):332–337. doi: 10.1038/s41586-019-1195-2. Epub 2019 May 1. Erratum in: Nature. 2019 Jun 17;: PMID: 31042697; PMCID: PMC6865822.

Melamed Z, López-Erauskin J, Baughn MW, Zhang O, Drenner K, Sun Y, Freyermuth F, McMahon MA, Beccari MS, Artates JW, et al. Premature polyadenylation-mediated loss of stathmin-2 is a hallmark of TDP-43-dependent neurodegeneration. Nat Neurosci. 2019 Feb;22(2):180–190. doi: 10.1038/s41593-018-0293-z. Epub 2019 Jan 14. PMID: 30643298; PMCID: PMC6348009.

Meneses A, Koga S, O’Leary J, Dickson DW, Bu G, Zhao N. TDP-43 Pathology in Alzheimer’s Disease. Mol Neurodegener. 2021 Dec 20;16(1):84. doi: 10.1186/s13024-021-00503-x. PMID: 34930382; PMCID: PMC8691026.

Morii H, Shiraishi-Yamaguchi Y, Mori N. SCG10, a microtubule destabilizing factor, stimulates the neurite outgrowth by modulating microtubule dynamics in rat hippocampal primary cultured neurons. J Neurobiol. 2006 Sep 1;66(10):1101–14. doi: 10.1002/neu.20295. PMID: 16838365.

Nag S, Yu L, Boyle PA, Leurgans SE, Bennett DA, Schneider JA. TDP-43 pathology in anterior temporal pole cortex in aging and Alzheimer’s disease. Acta Neuropathol Commun. 2018;6(1):33. Published 2018 May 1. doi:10.1186/s40478-018-0531-3

Ozon S, Maucuer A, Sobel A. The stathmin family -- molecular and biological characterization of novel mammalian proteins expressed in the nervous system. Eur J Biochem. 1997 Sep 15;248(3):794–806. doi: 10.1111/j.1432-1033.1997.t01-2-00794.x. PMID: 9342231.

Ozon S, Byk T, Sobel A. SCLIP: a novel SCG10-like protein of the stathmin family expressed in the nervous system. J Neurochem. 1998 Jun;70(6):2386–96. doi: 10.1046/j.1471-4159.1998.70062386.x. PMID: 9603203.

Prasad A, Bharathi V, Sivalingam V, Girdhar A, Patel BK. Molecular Mechanisms of TDP-43 Misfolding and Pathology in Amyotrophic Lateral Sclerosis. Front Mol Neurosci. 2019 Feb 14;12:25. doi: 10.3389/fnmol.2019.00025. PMID: 30837838; PMCID: PMC6382748.

Prudencio M, Humphrey J, Pickles S, Brown AL, Hill SE, Kachergus JM, Shi J, Heckman MG, Spiegel MR, Cook C, et al. Truncated stathmin-2 is a marker of TDP-43 pathology in frontotemporal dementia. J Clin Invest. 2020 Nov 2;130(11):6080–6092. doi: 10.1172/JCI139741. PMID: 32790644; PMCID: PMC7598060.

Riancho J, Paz-Fajardo L, López de Munaín A. Clinical and preclinical evidence of somatosensory involvement in amyotrophic lateral sclerosis. Br J Pharmacol. 2021 Mar;178(6):1257–1268. doi: 10.1111/bph.15202. Epub 2020 Aug 5. PMID: 32673410.

Riederer BM, Pellier V, Antonsson B, Di Paolo G, Stimpson SA, Lütjens R, Catsicas S, Grenningloh G. Regulation of microtubule dynamics by the neuronal growth-associated protein SCG10. Proc Natl Acad Sci U S A. 1997 Jan 21;94(2):741–5. doi: 10.1073/pnas.94.2.741. PMID: 9012855; PMCID: PMC19584.

Sanjana NE, Shalem O, Zhang F. Improved vectors and genome-wide libraries for CRISPR screening. Nat Methods. 2014 Aug;11(8):783–784. doi: 10.1038/nmeth.3047. PMID: 25075903; PMCID: PMC4486245.

Sasaki Y, Nakagawa T, Mao X, DiAntonio A, Milbrandt J. NMNAT1 inhibits axon degeneration via blockade of SARM1-mediated NAD+ depletion. Elife. 2016 Oct 13;5:e19749. doi: 10.7554/eLife.19749. PMID: 27735788; PMCID: PMC5063586.

Sasaki Y, Hackett AR, Kim S, Strickland A, & Milbrandt, J. Dysregulation of NAD+ Metabolism Induces a Schwann Cell Dedifferentiation Program. J. Neurosci. 38, 6546–6562 (2018).

Sasaki Y, Engber TM, Hughes RO, Figley MD, Wu T, Bosanac T, Devraj R, Milbrandt J, Krauss R, DiAntonio A. cADPR is a gene dosage-sensitive biomarker of SARM1 activity in healthy, compromised, and degenerating axons. Exp Neurol. 2020 Jul;329:113252. doi: 10.1016/j.expneurol.2020.113252. Epub 2020 Feb 19. PMID: 32087251; PMCID: PMC7302925.

Sentmanat MF, White JM, Kouranova E, Cui X. Highly reliable creation of floxed alleles by electroporating single-cell embryos. BMC Biol. 2022 Feb 4;20(1):31. doi: 10.1186/s12915-021-01223-w. PMID: 35115009; PMCID: PMC8815186.

Sharma A, Lyashchenko AK, Lu L, Nasrabady SE, Elmaleh M, Mendelsohn M, Nemes A, Tapia JC, Mentis GZ, Shneider NA. ALS-associated mutant FUS induces selective motor neuron degeneration through toxic gain of function. Nat Commun. 2016 Feb 4;7:10465. doi: 10.1038/ncomms10465. PMID: 26842965; PMCID: PMC4742863.

Shin JE, Miller BR, Babetto E, Cho Y, Sasaki Y, Qayum S, Russler EV, Cavalli V, Milbrandt J, DiAntonio A. SCG10 is a JNK target in the axonal degeneration pathway. Proc Natl Acad Sci U S A. 2012 Dec 26;109(52):E3696–705. doi: 10.1073/pnas.1216204109. Epub 2012 Nov 27. PMID: 23188802; PMCID: PMC3535671.

Shin JE, Geisler S, DiAntonio A. Dynamic regulation of SCG10 in regenerating axons after injury. Exp Neurol. 2014 Feb;252:1–11. doi: 10.1016/j.expneurol.2013.11.007. Epub 2013 Nov 15. PMID: 24246279; PMCID: PMC3947015.

Spiller KJ, Cheung CJ, Restrepo CR, Kwong LK, Stieber AM, Trojanowski JQ, Lee VM. Selective Motor Neuron Resistance and Recovery in a New Inducible Mouse Model of TDP-43 Proteinopathy. J Neurosci. 2016 Jul 20;36(29):7707–17. doi: 10.1523/JNEUROSCI.1457-16.2016. PMID: 27445147; PMCID: PMC6705561.

Stein R, Mori N, Matthews K, Lo LC, Anderson DJ. The NGF-inducible SCG10 mRNA encodes a novel membrane-bound protein present in growth cones and abundant in developing neurons. Neuron. 1988 Aug;1(6):463–76. doi: 10.1016/0896-6273(88)90177-8. PMID: 3272176.

Summers DW, Milbrandt J, DiAntonio A. Palmitoylation enables MAPK-dependent proteostasis of axon survival factors. Proc Natl Acad Sci U S A. 2018 Sep 11;115(37):E8746–E8754. doi: 10.1073/pnas.1806933115. Epub 2018 Aug 27. PMID: 30150401; PMCID: PMC6140512.

Summers DW, Frey E, Walker LJ, Milbrandt J, DiAntonio A. DLK Activation Synergizes with Mitochondrial Dysfunction to Downregulate Axon Survival Factors and Promote SARM1-Dependent Axon Degeneration. Mol Neurobiol. 2020 Feb;57(2):1146–1158. doi: 10.1007/s12035-019-01796-2. Epub 2019 Nov 7. PMID: 31696428; PMCID: PMC7035184.

Theunissen F, Anderton RS, Mastaglia FL, Flynn LL, Winter SJ, James I, Bedlack R, Hodgetts S, Fletcher S, Wilton SD, et al. Novel STMN2 Variant Linked to Amyotrophic Lateral Sclerosis Risk and Clinical Phenotype. Front Aging Neurosci. 2021 Mar 26;13:658226. doi: 10.3389/fnagi.2021.658226. PMID: 33841129; PMCID: PMC8033025.

van Es MA, Hardiman O, Chio A, Al-Chalabi A, Pasterkamp RJ, Veldink JH, van den Berg LH. Amyotrophic lateral sclerosis. Lancet. 2017 Nov 4;390(10107):2084–2098. doi: 10.1016/S0140-6736(17)31287-4. Epub 2017 May 25. PMID: 28552366.

Walker LJ, Summers DW, Sasaki Y, Brace EJ, Milbrandt J, DiAntonio A. MAPK signaling promotes axonal degeneration by speeding the turnover of the axonal maintenance factor NMNAT2. Elife. 2017 Jan 17;6:e22540. doi: 10.7554/eLife.22540. PMID: 28095293; PMCID: PMC5241118.

Wang Q, Zhang Y, Wang M, Song WM, Shen Q, McKenzie A, Choi I, Zhou X, Pan PY, Yue Z, Zhang B. The landscape of multiscale transcriptomic networks and key regulators in Parkinson’s disease. Nat Commun. 2019 Nov 20;10(1):5234. doi: 10.1038/s41467-019-13144-y. PMID: 31748532; PMCID: PMC6868244.

Wilson RS, Yu L, Trojanowski JQ, et al. TDP-43 pathology, cognitive decline, and dementia in old age. JAMA Neurol. 2013;70(11):1418–1424. doi:10.1001/jamaneurol.2013.3961

