## Supplementary figures and images for "Loss of Stathmin-2, a hallmark of TDP-43-associated ALS, causes motor neuropathy"

### Supplementary Fig. 1

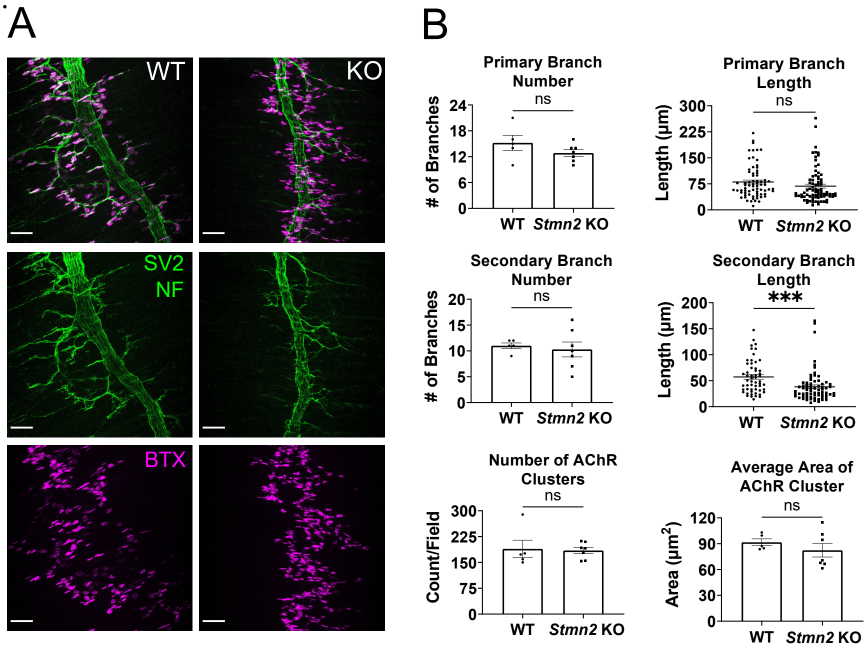

### Supplementary Fig. 2

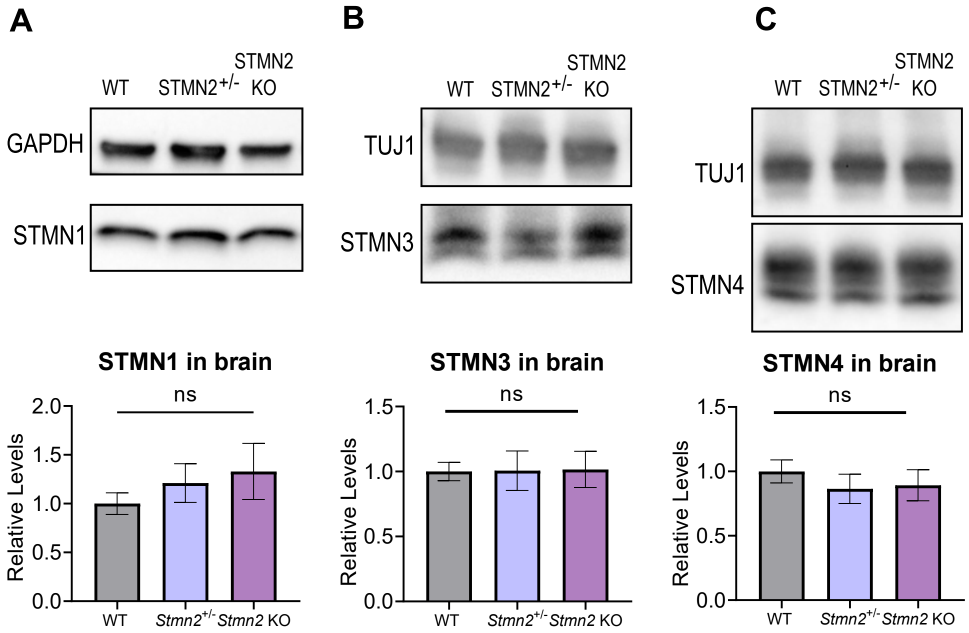

### Supplementary Fig. 3

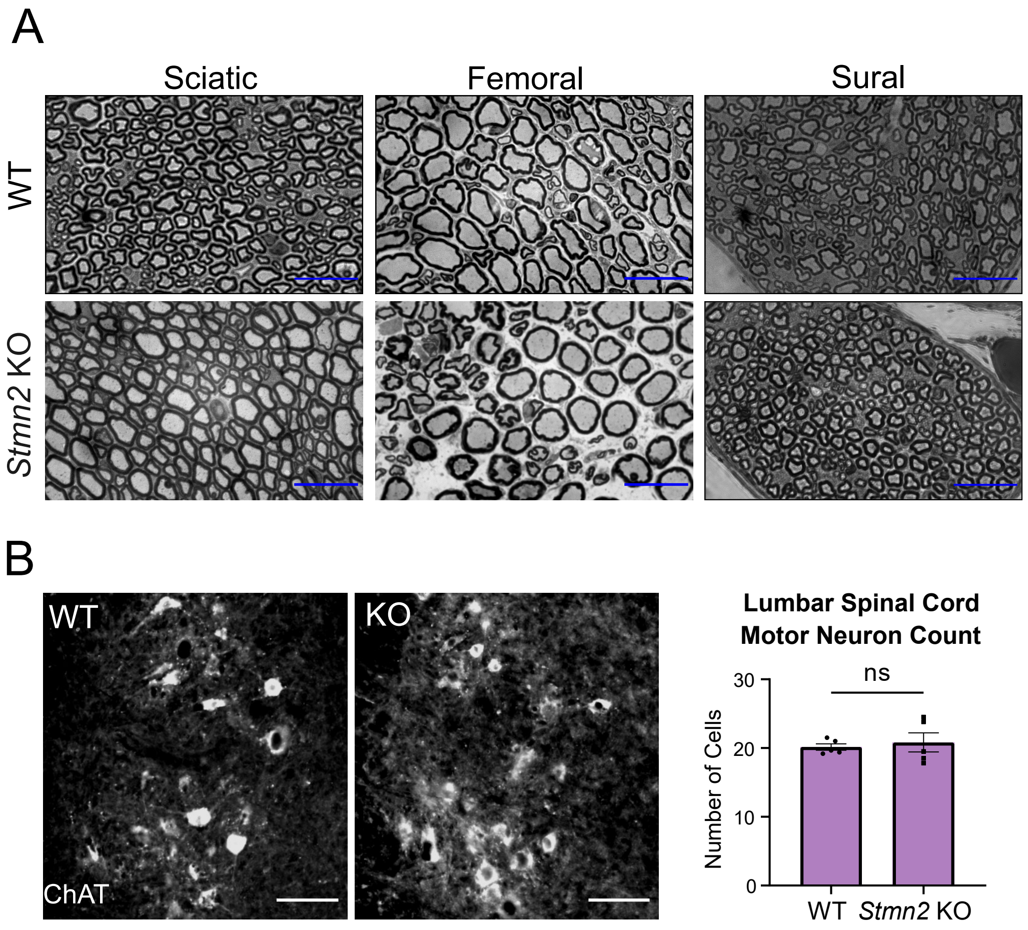

### Supplementary Fig. 4

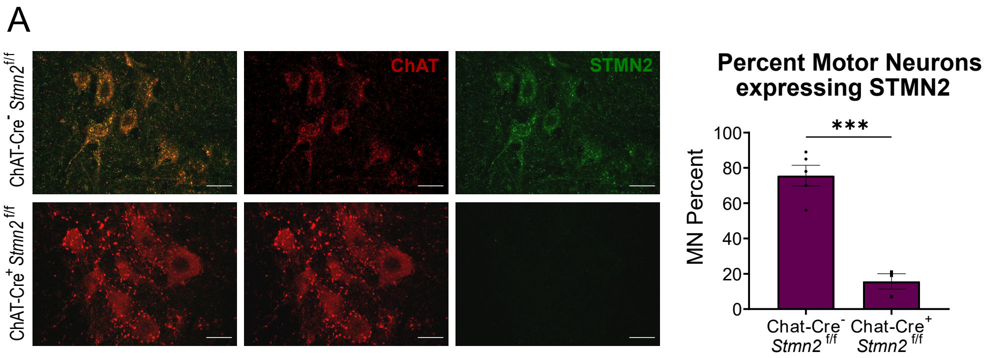

### Supplementary Fig. 5

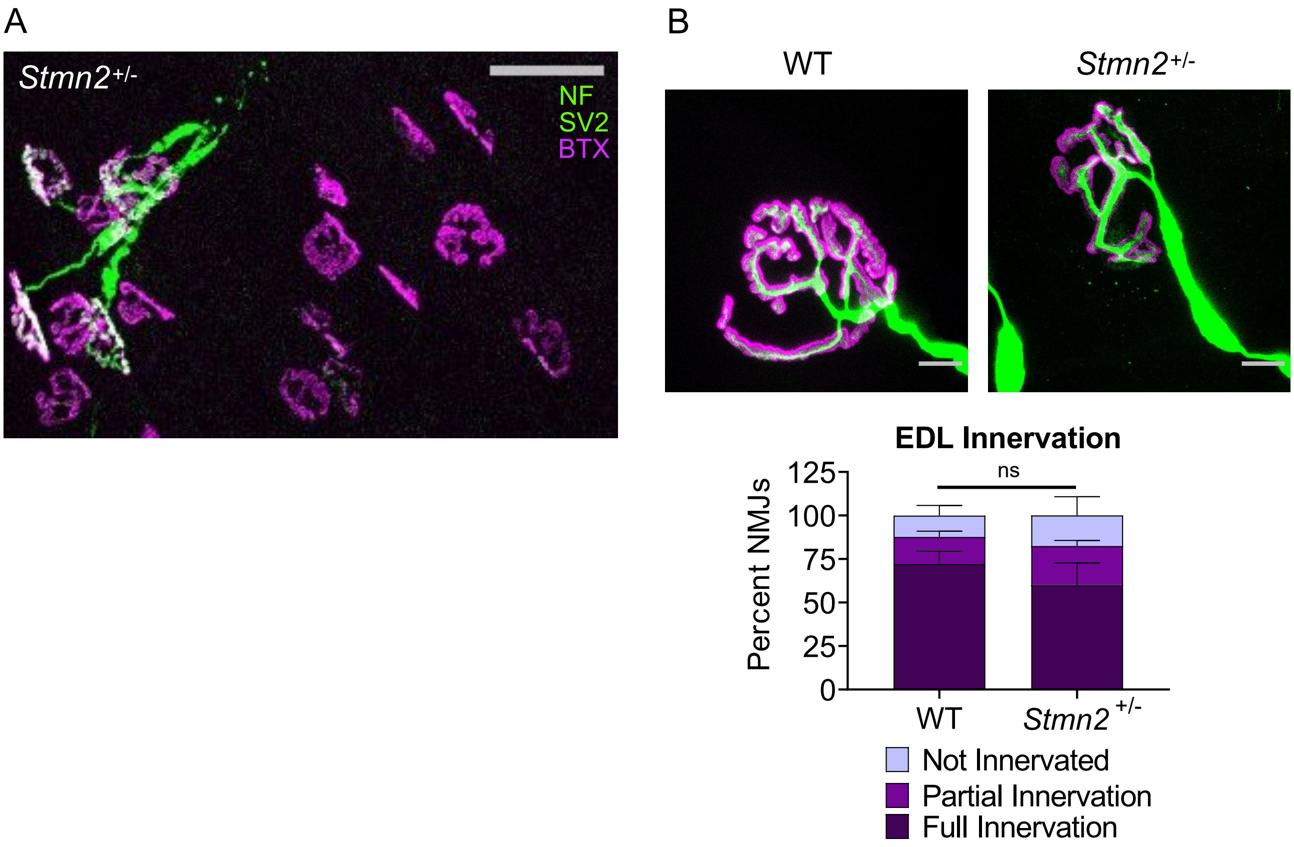
